# Extracellular condensates (ECs) are endogenous modulators of HIV transcription and latency reactivation

**DOI:** 10.1101/2024.09.14.613037

**Authors:** Wasifa Naushad, Lakmini S Premadasa, Bryson C. Okeoma, Mahesh Mohan, Chioma M. Okeoma

## Abstract

Persistence of human immunodeficiency virus (HIV) latent reservoir is the major challenge to HIV cure because the latent reservoir is not eliminated by antiretroviral therapy (ART), and they serve as sources for viral rebound upon cessation of ART. Mechanisms regulating viral persistence are not well understood. This study used model systems of post-integration latency to explore the role of basal ganglia (BG) isolated extracellular condensates (ECs) in reprogramming HIV latent cells. We found that BG ECs from uninfected macaques (VEH) and SIV infected macaques (VEH|SIV) activate latent HIV transcription in various model systems. VEH and VEH|SIV ECs significantly increased expression of viral antigen in latently infected cells. Activation of viral transcription, antigen expression, and latency reactivation was inhibited by ECs from the brain of macaques treated with Delta-9-tetrahydrocannabinol (THC) and infected with SIV (THC|SIV). Virus produced by latently infected cells treated with VEH|SIV ECs potentiated cell-cell and cell-free HIV transmission. VEH|SIV ECs also reversed dexamethasone-mediated inhibition of HIV transcription while TNFα-mediated reactivation of latency was reversed by THC|SIV ECs. Transcriptome and secretome analyses of total RNA and supernatants from latently infected cells treated with ECs revealed significant alteration in gene expression and cytokine secretion. THC|SIV ECs increased secretion of Th2 and decreased secretion of proinflammatory cytokines. Most strikingly, while VEH/SIV ECs robustly induced HIV RNA in latently HIV-infected cells, long-term low-dose THC administration enriched ECs for anti-inflammatory cargo that significantly diminished their ability to reactivate latent HIV, an indication that ECs are endogenous host factors that may regulate HIV persistence.

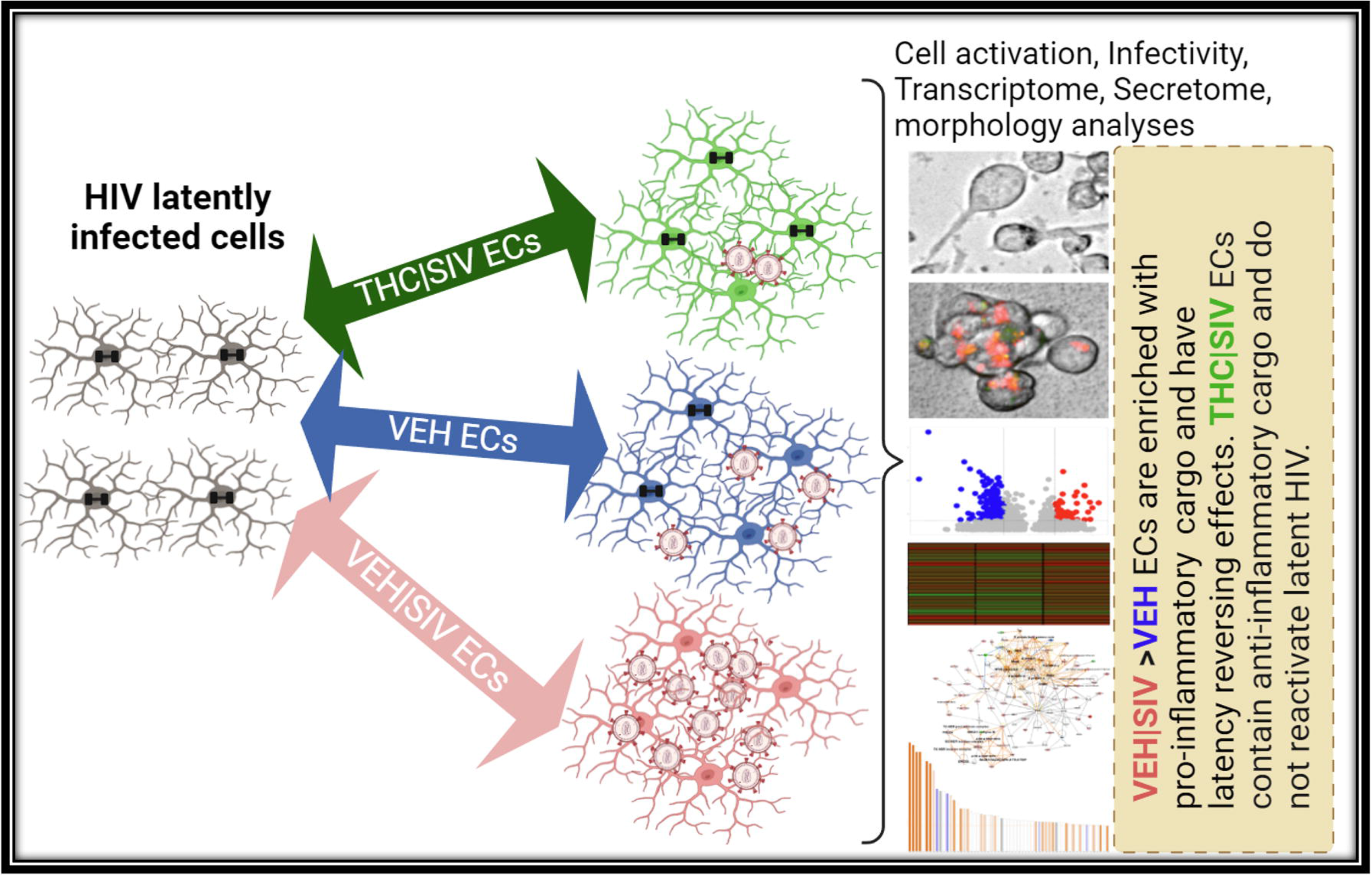

**Highlights:**

- ECs isolated from SIV infected macaques (VEH|SIV ECs) is a positive regulator of LTR-dependent HIV transcription and production of infectious viral particles in vitro.
- ECs isolated from THC treated SIV infected macaques (THC|SIV ECs) prevents the transcription and reactivation of HIV in latently infected cells and prevents production of viral particles in vitro.
- ECs reprogram host transcriptome and secretome in manners that or suppress promote reactivation of latent HIV reservoir.

The above highlights led to the conclusion that while VEH/SIV ECs robustly induced HIV RNA in latently HIV-infected cells, long-term low-dose THC administration enriched ECs for anti-inflammatory cargo that significantly diminished their ability to reactivate latent HIV.

## Introduction

The cure for HIV requires rigorous assessment of tissue reservoirs and the extracellular host factors that regulate HIV persistence despite ART. The central nervous system (CNS) is a unique site where HIV persists. Circulating CD4 T cells are the major HIV reservoir in the periphery, and these cells migrate into the CNS and may contribute to HIV persistence in the brain [1]. Additionally, myeloid cells (monocytes, macrophages, and microglia) are also HIV targets/reservoirs. HIV may persist in brain myeloid cells [2-6] and these cells may contribute to the size of the latent reservoir, as revealed by the presence of HIV DNA in postmortem brains from people with HIV (PWH) [7, 8]. However, the host factors that regulate viral persistence in the CNS are not well understood. Thus, identifying the endogenous parameters that contribute to HIV persistence is of significant interest. We predict that extracellular condensates (ECs) may function as endogenous regulators of HIV persistence.

ECs are non-lipid carriers of protein and nucleic acid cargo [9, 10] that may act in a paracrine manner to mediate cell to cell interaction, modulate host responses, and remodel cellular biological processes. Of particular interest and a never before studied topic is the link between endogenous secreted factors, such as ECs and their ability to reactivate latent HIV, especially given that they can be modified by the microenvironment, can be generated in vivo, and can trigger complex cellular response. Indeed, it has been suggested that extracellular vesicles in the brain may play a role in HIV pathogenesis [11].

Cannabis use is high among PWH and was associated with reduced immune activation [12], lower plasma HIV RNA [13] and most strikingly, significantly reduced proviral HIV DNA burden in multiple tissues [14]. Recently, our group showed that long-term low dose delta-9-tetrahydrocannabinol (THC) administration reduced neuroinflammation[15] and stimulated the release of blood EVs that induced divergent structural adaptations and signaling cues in chronically SIV-infected rhesus macaques [16]. We further showed that the basal ganglia (BG) contains ECs and EVs and that BG EVs serve as a vehicle with the potential to disseminate SIV and THC induced changes within the CNS [11]. Since the BG [17, 18] is a major HIV target/reservoir, these findings collectively underscore the need for further studies to investigate the effects of BG ECs and EVs on HIV persistence in the context of THC use.

The current study combined the SIVmac251-infected rhesus macaque model of HIV infection and pathogenesis, BG ECs, and in vitro model systems of post-integration latency to investigate the effect of ECs on reactivation of latent HIV. We found specific effects of ECs in reactivating latent HIV in J-Lat -GFP, J-Lat Tat-GFP, U1 monocytic, and hμglia (HC69) microglia cells. Moreover, we demonstrated EC-mediated alteration in transcriptome and secretome of HIV latently infected cells that were distinct in cells treated with VEH|SIV versus THC|SIV ECs. The transcriptome and secretome changes culminated in alteration of important pathways, especially the NF-κB, replication factor C4 (RFC4), and CDK complexes, that are important regulators of HIV transcription, and reactivation of latent HIV. Altogether, our results demonstrate ECs as endogenous regulators of HIV latency with potential to control the size of viral reservoirs and persistence.

## Results

### Assessment of plasma and BG viral loads

**Brain** viral loads in the two groups ranged from 0.01 to 2.0 10^6^/mg (p=0.100) RNA (**Table 1**) and were partly published previously [11]. ECs were isolated from BG tissues of VEH (n=3), VEH|SIV (n=4), THC|SIV (n=4) RMs (**Table 1)**. VEH represents the uninfected control (n=3). All THC/SIV RMs initiated twice daily THC injections (i/m) at 0.18 mg/kg two weeks before SIV and continued thereafter at 0.32 mg/kg until necropsy (**Supplemental material**).

**Table 1.** Animal IDs, SIV inoculum, duration of infection, viral loads and brain histopathology in vehicle or delta-9-tetrahydrocannabinol (Δ^9^-THC) treated chronic SIV-infected and uninfected rhesus macaques

| Animal ID | SIV Inoculum | Duration of Infection | Plasma viral loads $10^6$/mL | Brain viral loads $10^6$/mg RNA | Brain Histopathology | Opportunistic Infections |
| --- | --- | --- | --- | --- | --- | --- |
| **Chronic SIV-Infected and Vehicle treated (Group 1)** |  |  |  |  |  |  |
| IV95 | SIVmac251 | 180 | 0.02 | 2.0 | ND | ND |
| JD66 | SIVmac251 | 180 | 0.04 | 0.2 | ND | ND |
| JR36 | SIVmac251 | 180 | 0.5 | 0.2 | ND | ND |
| JH47 | SIVmac251 | 180 | 2 | 0.07 | ND | ND |
| JI45 | SIVmac251 | 180 | 3 | 0.01 | ND | ND |
| JT80 | SIVmac251 | 180 | 1 | 0.04 | ND | ND |
| JC85 | SIVmac251 | 180 | 0.02 | 0.09 | ND | ND |
| IV90 | SIVmac251 | 180 | 0.02 | 0.06 | ND | ND |
| **Uninfected Controls (Group 3)** |  |  |  |  |  |  |
| IR97 | NA | NA | NA | NA | NA | NA |
| IT18 | NA | NA | NA | NA | NA | NA |
| GT18 | NA | NA | NA | NA | NA | NA |

NA- Not applicable, ND- None detected

### VEH|SIV BG ECs reactivate HIV latency in J Lat T cells

J-Lat GFP cells (**Supplemental Fig. 1A, 1B**) were treated with DiR-labeled PBS, or VEH, VEH|SIV, THC|SIV ECs (**Table 1**). Internalization of ECs (**Fig. 1A**) and GFP expression as an evidence of HIV reactivation (**Figs. 1B, 1C**) were analyzed at 24 h. Microscopic data showed that VEH|SIV ECs significantly increased GFP expression in J-Lat GFP cells, which was confirmed by flow cytometry with 3.8-fold change in GFP expression in VEH|SIV-treated cells compared to PBS treated cells (**Fig. 1D**). The effect of ECs on latent HIV reactivation was validated with J-Lat-Tat-GFP (**Figs. 1E-1H**).

**Figure 1.**
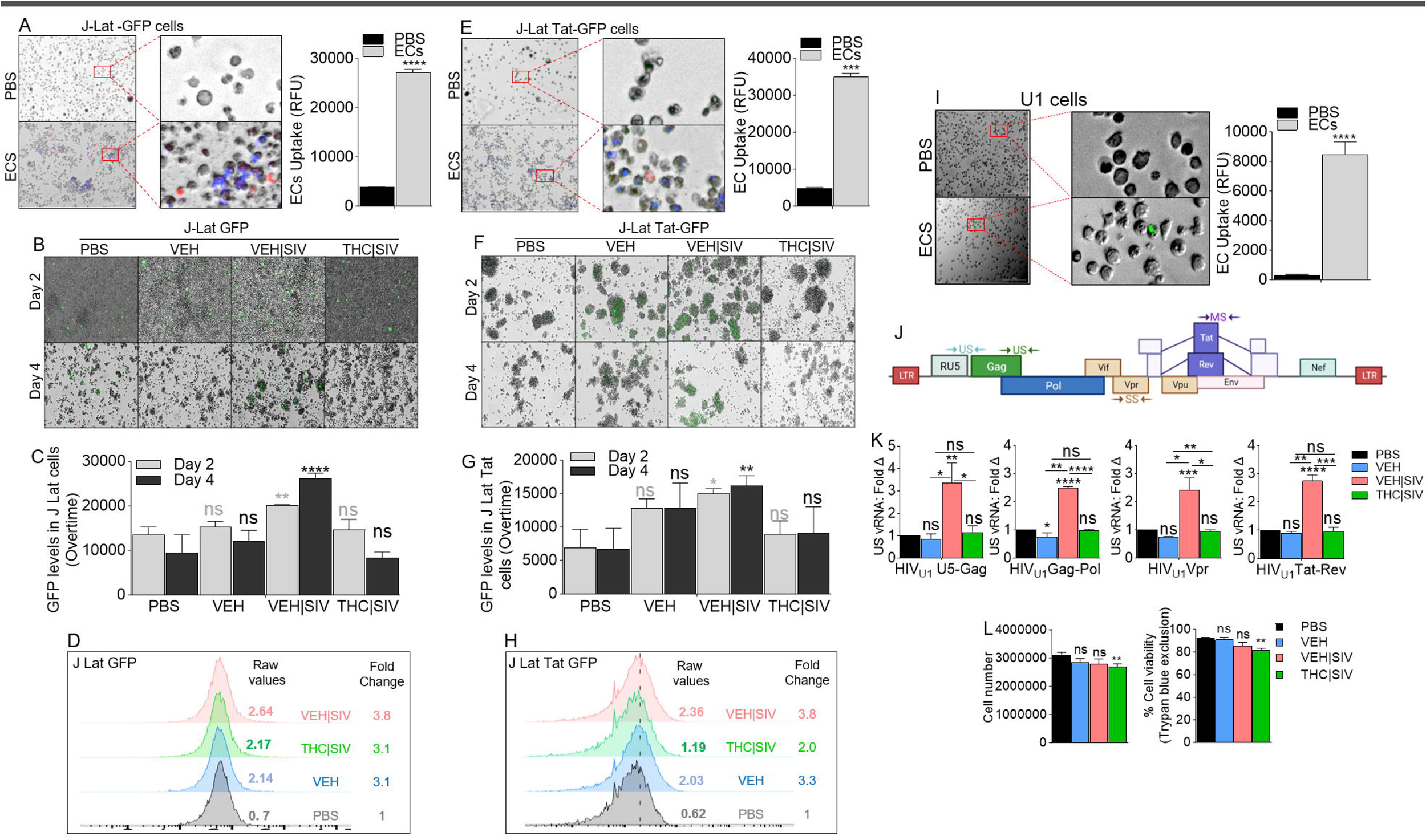
Basal ganglia derived ECs activate HIV latently infected cells. **A**) Representative microscopic images of DIR-labeled ECs internalized by J-Lat -GFP cells (left) and the level of internalized ECs (bars, right). **B**) Representative microscopic images of EC-treated J-Lat -GFP cells showing HIV reactivation (GFP expression, green) after 4 days of treatment. Representative data from day 4 treatment presented. **C**) The level of HIV reactivation analyzed by microscopy was quantified on days 2 and 4. **D**) Flow cytometry analysis of HIV reactivation (GFP expression, green) after 4 days of treatment. Numbers indicate raw values (left) or fold change (treatment/PBS) (right). **E**) Microscopic image of internalized DIR-labeled ECs by J-Lat Tat-GFP cells (left) and the level of internalized ECs (bars, right). **F**) Microscopic images of EC-treated J-Lat -GFP cells showing HIV reactivation (GFP expression, green) after 4 days of treatment. Representative data from day 4 treatment presented. **G**) The level of HIV reactivation analyzed by microscopy was quantified on days 2 and 4. **H**) Flow cytometry analysis of HIV reactivation (GFP expression, green) after 4 days of treatment. Numbers indicate raw values (left) or fold change (treatment/PBS) (right). **I**) Microscopic image of internalized SYTO RNASelect green fluorescent-labeled U1 cells (left) and the level of internalized ECs (bars, right). **J**) HIV genome showing the different HIV genes. Arrows indicate the positions of forward and reverse primers used in the analysis of HIV transcription. **K**) RT-qPCR analysis of the levels of intracellular viral RNA. Four HIV (unspliced [U5-Gag, Gag-Pol], singly spliced [Vpr], and multiply spliced [Tat-Rev]) transcripts were measured. **L**) Analysis of cell density following 4-day treatment with EC as assessed by cell number (left) and trypan blue exclusion assay (right). All experiments were repeated three times. Statistical differences were assessed by ordinary one-way ANOVA with Tukey’s correction and by Binary Student’s t tests (Welch’s correction). **** p < 0.001, *** p < 0.005, ** p < 0.01, p < 0.05, and ns = non-significant.

### VEH|SIV but not THC|SIV BG ECs reverse HIV-1 latency in U1 cells

HIV-1 latency reversal potential of VEH|SIV BG ECs was assessed using U1 monocytes as an in vitro model of postintegration latency [19][20] (**Supplemental Fig. 1A, 1C**). ECs were internalized by U1 cells 24 h after treatment (**Fig. 1I**). The degree to which HIV expression is activated by ECs at different stages of HIV transcription was assessed using a panel of primers that amplify unspliced [U5-Gag, Gag-Pol], singly spliced [Vpr], and multiply spliced [Tat-Rev] transcripts (**Table 2, Fig. 1J**). Compared to PBS, VEH, and THC|SIV ECs, VEH|SIV ECs reversed HIV-1 latency as shown by the significant increases in all intracellular HIV RNA types (**Fig. 1K**). Since cell death in reactivated cells is a key mechanism to reduce HIV reservoir cells in vivo [21, 22], we showed that 24 h after treatment, THC|SIV but not VEH and VEH|SIV ECs modestly but significantly reduced U1 cell numbers (**Fig. 1L left**) and viability (**Fig. 1L right**).

### VEH|SIV but not THC|SIV BG ECs promote production of infectious HIV

Compared to PBS and THC|SIV ECs, VEH and VEH|SIV ECs induced significant expression of p24 and p17 in U1 cells (**Fig. 2A, B**). Cell-free HIV particles from U1 cells treated for 96 h with PBS or ECs were assessed for extracellular reverse transcriptase (exRT) as a proxy for the concentration of cell-free virus [23-25]. Compared to PBS and VEH ECs that did not change exRT, THC|SIV ECs significantly decreased while VEH|SIV ECs significancy increased exRT induction (**Fig. 2C**). HIV particles produced in the presence of the VEH|SIV but not VEH and THC|SIV ECs were infectious as shown by GFP expression in TZM-GFP indicator cells on days 1 (**Figs. 2D, E**) and 2 (**Figs. 2F, G**).

**Figure 2.**
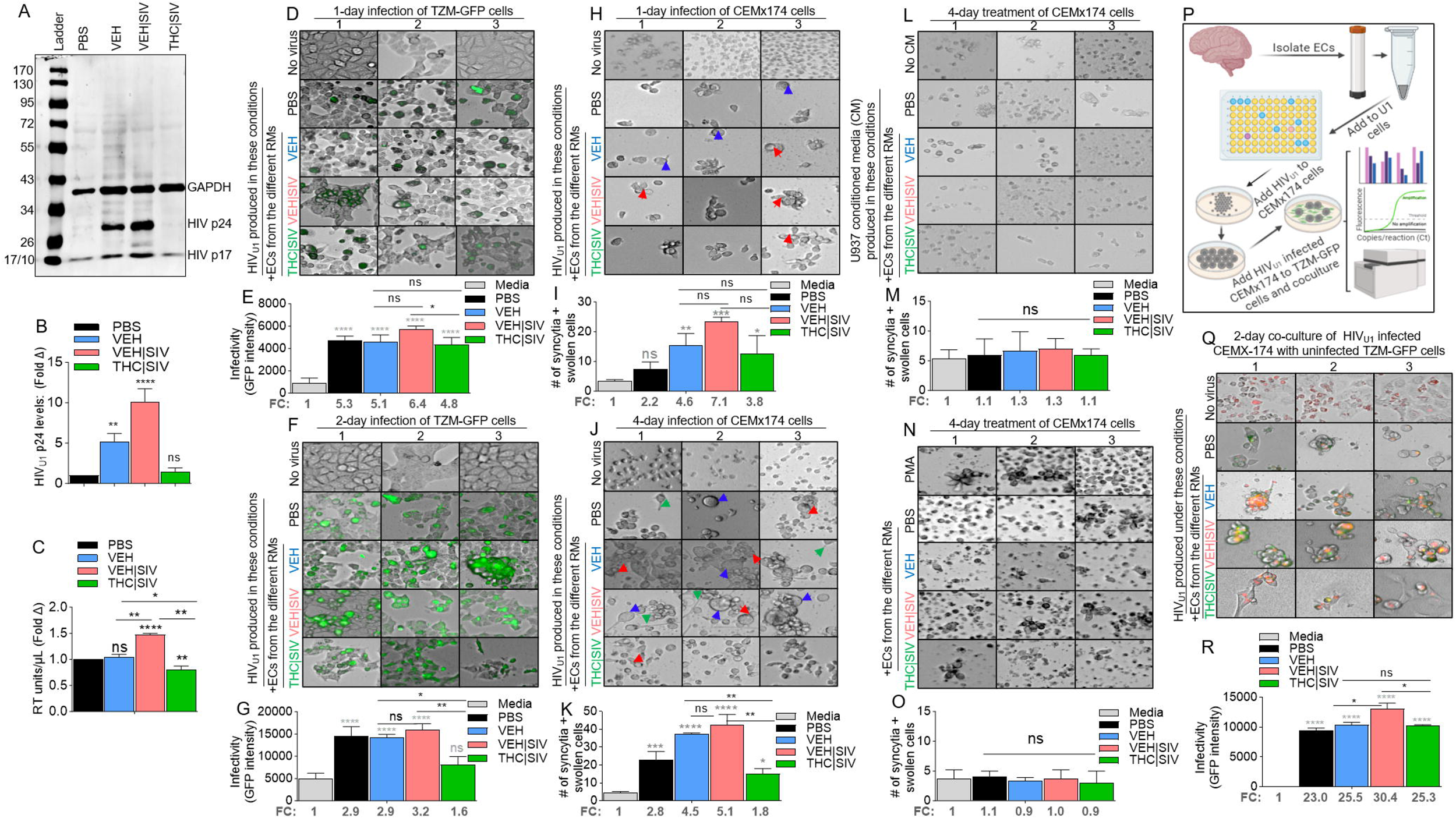
Potency of HIV latency reversal by VEH < VEH|SIV ECs and THC|SIV ECs. **A**) Representative western blot image of intracellular expression of HIV antigens p24 and p17, with host GAPDH used as loading control. **B**) The levels of HIV p24 expression analyzed by densitometry (ImageJ) and presented as fold change (treatment/PBS). **C**) The level of extracellular HIV RT released by EC-treated U1 cells into the culture supernatants after 4 days of treatment. **D-G**) Clarified U1 cell supernatants collected on day 4 of treatment were added to the indicator cells – TZM-GFP cells and cultured for (**D, E**) one day or (**F, G**) two days. **D, F**) Representative microscopic images of HIV infection (GFP expression, green) after 1 and 2 days of infection respectively. **E, G**) Quantification of HIV infection (GFP expression, green) after 1 and 2 days of infection respectively. The numbers below the bars indicate fold change (treatment/PBS). “No virus” panel shows the level of autofluorescence detected by the instrument. **H-K**) The same clarified U1 cell supernatants collected on day 4 of treatment were added to CEMx174 cells and cultured for (**H, I**) one day or (**J, K**) four days. **H, J**) Representative microscopic images of cell morphology. The number of syncytia and large cells were quantified on **I**) day 1 and **K**) day 4. Red, blue, and green arrows denote syncytia, large cells, and filopodia-like protrusions respectively. **L-M**) Clarified supernatants collected on day 4 from EC-treated U937 cells were added to CEMx174 cells and cultured for 4 days. **L**) Representative microscopic images of cell morphology. **M**) The number of syncytia and large cells were quantified. **N-O**) CEMx174 cells were treated with VEH, VEH|SIV, THC|SIV ECs for 4 days. No virus = cells treated with PBS but not media from EC-treated U1 cells. No CM = cells treated with regular media but not conditioned media. **N**) Microscopic image of cell morphology. **M**) The number of syncytia and large cells were quantified. **P**) Schematic of HIV transfer assay from CEMx174 cells infected with EC-induced HIV cocultured with uninfected indicator TZM-GFP cells. **Q**) Representative microscopic images of HIV transfer assay from CEMx174 (red) to TZM-GFP cells. Green and orange (green + red) indicate infection. **R**) The level of infection (GFP expression). The numbers below the bars indicate fold change (treatment/PBS). All experiments were repeated three times. Statistical differences were assessed by ordinary one-way ANOVA with Tukey’s correction and by Binary Student’s t tests (Welch’s correction). **** p < 0.001, *** p < 0.005, ** p < 0.01, p < 0.05, and ns = non-significant.

### VEH|SIV but not THC|SIV BG ECs promote HIV-induced syncytia formation

Compared to CEMx174 cells treated with PBS or THC|SIV ECs, higher levels of cell-cell fusion were observed in cells treated with VEH|SIV > VEH ECs on days 1 (**Figs. 2H, I**) and 4 (**Figs. 2J, K**). The appearance of other morphological changes, including swollen cells (**Fig. 2J, blue arrows**) and filopodia-like protrusions (**Fig. 2J, green arrows**) were observed. Interestingly, supernatants from HIV-negative U937 cells (parental cells of U1) treated with PBS or ECs did not induce syncytia (**Figs. 2L, M**) and direct addition of PMA or any of the ECs to CEM x174 cells did not induce syncytia formation (**Figs. 2N, O**). These data indicate that syncytia promoting ability of ECs requires the presence of HIV.

### VEH|SIV but not THC|SIV BG ECs potentiate cell-to-cell HIV transfer

Coculture of HIV infected CEM x174 cells with uninfected TZM-GFP cells (**Fig. 2P**) resulted in virus transfer to the TZM-GFP cells. The level of virus transfer depends on the source of ECs used to reactivate U1 virus for infection. Thus, cells infected with virus from VEH|SIV ECs mediated the highest level of HIV transfer (**Figs. 2Q, 2R**). GFP expression in uninfected TZM-bl cells cocultured with infected CEM x174 cells suggest that the EC-reactivated latent HIV virus is fully replicative, and virus dissemination may have occurred through cell-cell contact, with the potential to mediate successful establishment of HIV in atypical cells.

### Durability of BG ECs as latency regulating agents

U1 cells treated with single or repeated doses of PBS or ECs (**Fig. 3A**) over time express HIV RNA mirroring the ECs. HIV RNA in cells treated with single dose of ECs returned to near baseline on day 16 (**Fig. 3B, left, Table 3**) in contrast with cells treated with continuous dose of ECs where HIV RNA continued to increase up to day 16 (**Fig. 3B, right, Table 3**). The trend of HIV RNA increase was VEH|SIV>VEH>THC|SIV=PBS.

**Table 3.**
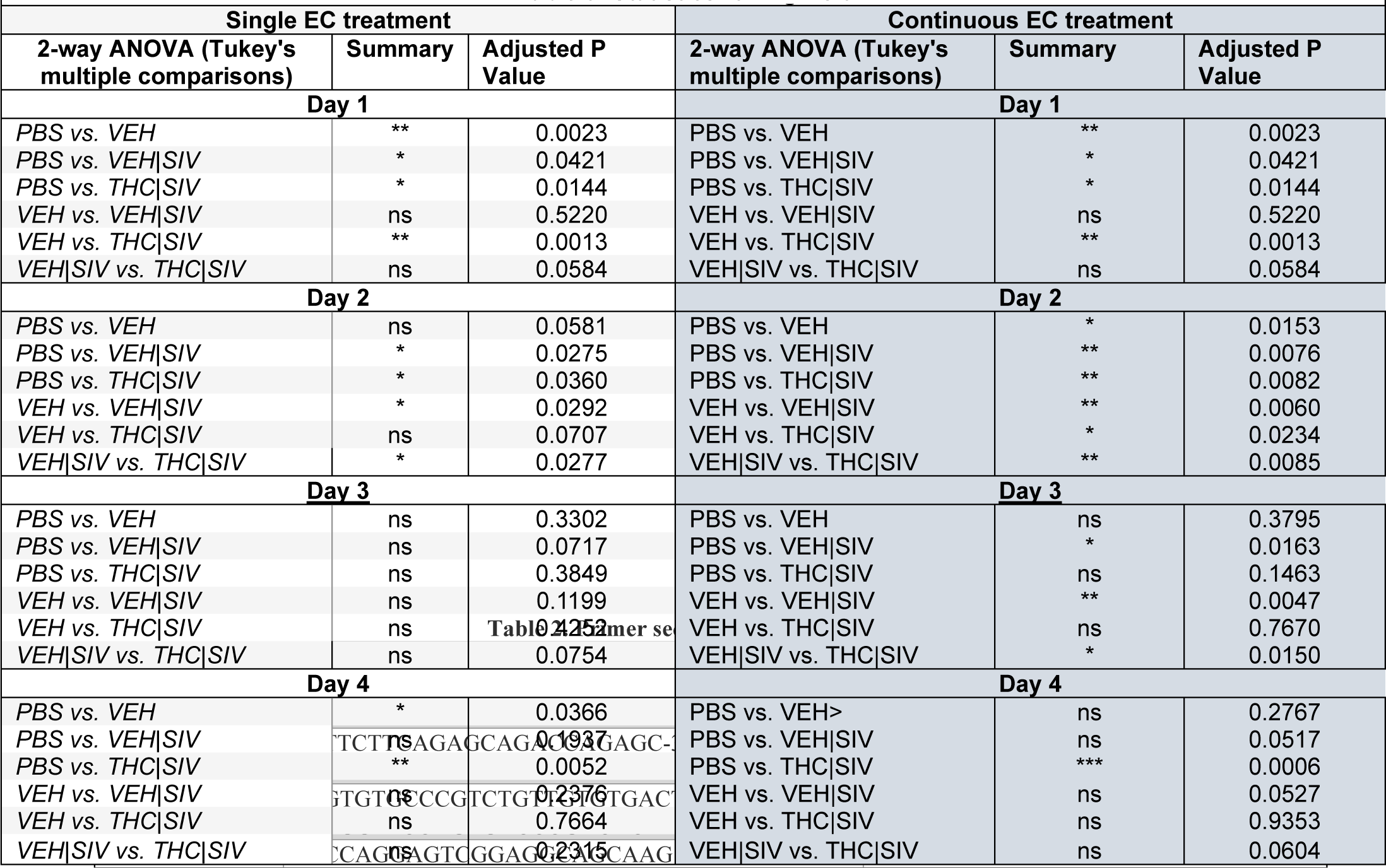
Statistics for Figure 3B.

**Figure 3.**
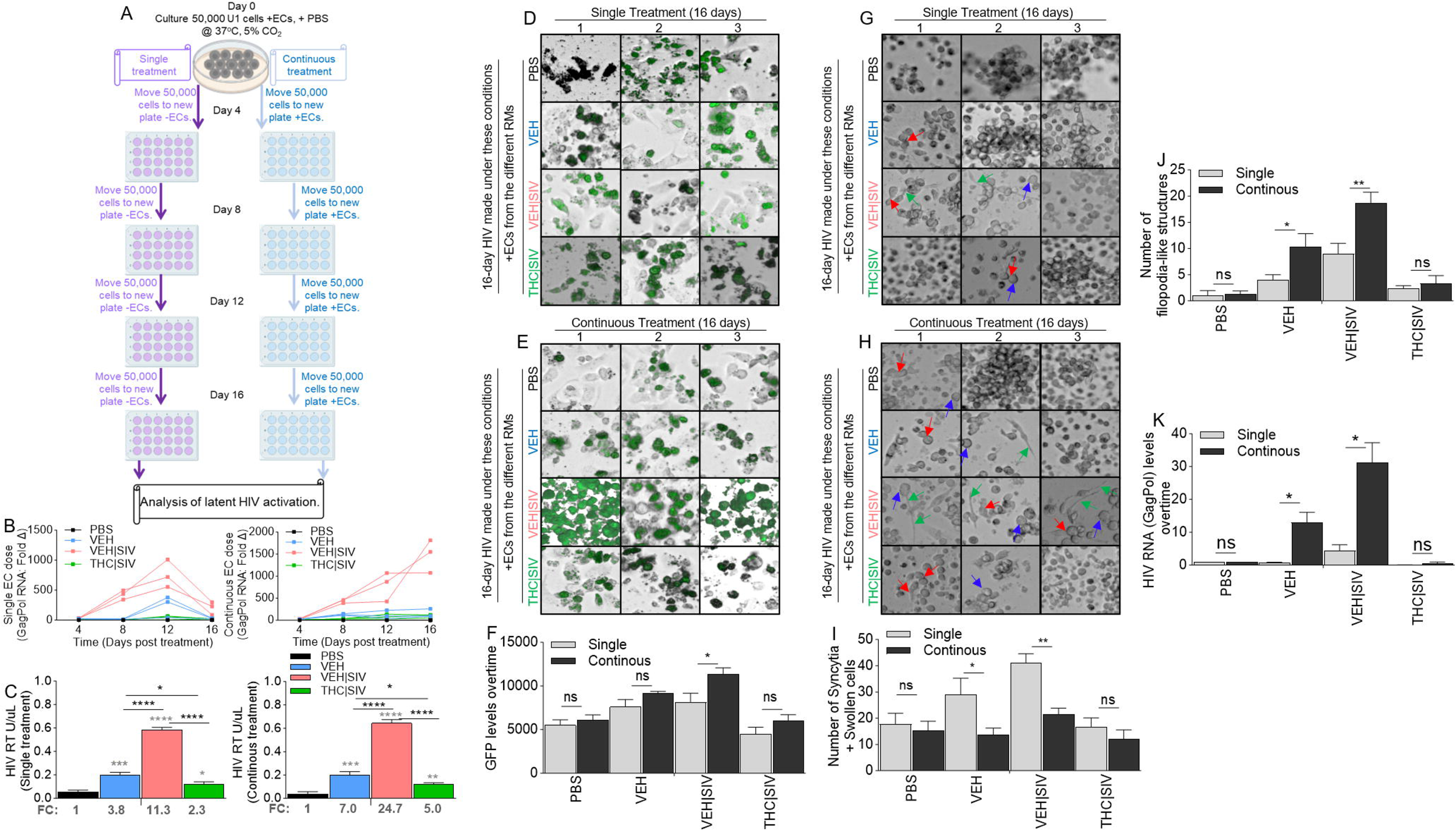
Basal ganglia derived EC-mediated activation or inhibition of latent HIV reactivation is durable. **A**) Schematic of single and continuous cell treatment with ECs indicating the number of cells, the time of cell passage, the time of EC treatment, and the length of cell culture. **B**) RT-qPCR analysis of the levels of intracellular HIV Gag-Pol expression for single EC treatment (left) and continuous EC treatment (right) of U1 cells. **C**) The level of extracellular HIV RT released by EC-treated U1 cells into the culture supernatants after single EC treatment (left) and continuous EC treatment (right). The numbers below the bars indicate fold change (treatment/PBS). **D-F**) Representative microscopic images of HIV infection (GFP expression, green) after **D**) single EC treatment and **E**) continuous EC treatment. **F**) Quantification of HIV infection (GFP expression, green) for single and continuous EC treatment. **G-H**) Representative microscopic images of cell morphology for **G**) single EC treatment and **H**) continuous EC treatment. Red, blue, and green arrows denote syncytia, large cells, cytoplasmic extension respectively. **I**) The number of syncytia and large. **J, K**) RT-qPCR analysis of the levels of intracellular HIV Gag-Pol expression in CEMx174 cells treated with supernatants from U1 cells subjected to single EC treatment (left) and continuous EC treatment (right). All experiments were repeated three times. Statistical differences were assessed by ordinary one-way ANOVA with Tukey’s correction and by Binary Student’s t tests (Welch’s correction). **** p < 0.001, *** p < 0.005, ** p < 0.01, * p < 0.05, and ns = non-significant. Statistical differences for Figure 3B were assessed by ordinary 2-way ANOVA with Tukey’s multiple comparison test as presented on **Table 3**.

Analysis of exRT on day 16 of single or continuous doses of ECs showed similar trends with HIV RNA (**Fig. 3C**). Importantly, continuous treatment doubled the amount of exRT (11.3 vs 24.7) in culture supernatant compared to single (**Fig. 3C**). Compared to single dose of ECs, continuous doses of ECs resulted in increased GFP expression in TZM-GFP cells (**Figs. 3D-F**). HIV infectivity was confirmed by syncytia in recipient CEMx174 cells. The number of syncytia and swollen cells were significantly higher in VEH|SIV > VEH cells that received single dose of ECs (**Figs. 3G, 3I**), compared to continuous doses of ECs (**Figs. 3H, 3I**). THC|SIV did not increase the number of syncytia compared to PBS (**Figs. 3G-3I**). Interestingly, the number of filopodia-like structures were significantly higher in VEH|SIV > VEH cells that received multiple doses of ECs (**Figs. 3H, 3J**) compared to single dose of ECs (**Figs. 3G, 3J**). The effects of ECs on the release of infectious virus were confirmed by analysis of viral RNA in recipient CEMx174 cells, where ECs induced significant HIV GagPol expression in the following order VEH|SIV > VEH > THC|SIV = PBS in that order (**Figs. 3K**).

### EH|SIV BG ECs reactivate latent HIV in microglia

To directly assess the effects of ECs as latency-reversing agents in microglia cells, we used hµglia -HC69 cells [26] that express GFP upon activation (**Supplemental Fig. 1A**). Cells treated with ECs for 24 h internalized ECs (**Fig. 4A**). VEH|SIV ECs reactivated latent HIV in hµglia cells similar to the level seen in TNFα (10 ng/mL) treated cells (**Figs. 4B, 4C**). Reactivation of latent HIV was progressive and significantly increased from day 1 to day 4 where VEH|SIV > VEH (**Figs. 4B, 4C, Table 4**). Hµglia cells treated with THC|SIV ECs had quiescent phenotype similar to what was observed in PBS or dexamethasone (DEX, 3 mM/mL) treated cells (**Figs. 4B, 4C, Table 4**).

**Table 4.** Statistics for Figure 4C.

| Table 4: Statistics for Figure 4C |  |  |  |  |  |
| --- | --- | --- | --- | --- | --- |
| 2-way ANOVA (Tukey's multiple comparisons) | Summary | Adjusted P Value | 2-way ANOVA (Tukey's multiple comparisons) | Summary | Adjusted P Value |
| Day 1 |  |  | Day 2 |  |  |
| PBS vs. DEX | ns | 0.4925 | PBS vs. DEX | ns | 0.7456 |
| PBS vs. TNFα | * | 0.0233 | PBS vs. TNFα | * | 0.0287 |
| PBS vs. VEH | ns | 0.8943 | PBS vs. VEH | ns | 0.2834 |
| PBS vs. VEH\|SIV | ns | 0.2488 | PBS vs. VEH\|SIV | * | 0.0114 |
| PBS vs. THC\|SIV | ns | 0.6960 | PBS vs. THC\|SIV | ns | 0.9575 |
| DEX vs. TNFα | * | 0.0227 | DEX vs. TNFα | * | 0.0163 |
| DEX vs. VEH | ns | 0.2514 | DEX vs. VEH | ns | 0.1456 |
| DEX vs. VEH\|SIV | ns | 0.1624 | DEX vs. VEH\|SIV | * | 0.0108 |
| DEX vs. THC\|SIV | ns | 0.9178 | DEX vs. THC\|SIV | ns | 0.4812 |
| TNFα vs. VEH | * | 0.0248 | TNFα vs. VEH | ns | 0.0600 |
| TNFα vs. VEH\|SIV | ns | 0.2063 | TNFα vs. VEH\|SIV | ns | 0.1585 |
| TNFα vs. THC\|SIV | * | 0.0324 | TNFα vs. THC\|SIV | * | 0.0292 |
| VEH vs. VEH\|SIV | ns | 0.3528 | VEH vs. VEH\|SIV | ns | 0.5047 |
| VEH vs. THC\|SIV | ns | 0.3632 | VEH vs. THC\|SIV | ns | 0.4295 |
| VEH\|SIV vs. THC\|SIV | ns | 0.1946 | VEH\|SIV vs. THC\|SIV | * | 0.0233 |
| Day 3 |  |  | Day 4 |  |  |
| PBS vs. DEX | ns | 0.4134 | PBS vs. DEX | ns | 0.2990 |
| PBS vs. TNFα | ns | 0.0645 | PBS vs. TNFα | * | 0.0318 |
| PBS vs. VEH | ** | 0.0034 | PBS vs. VEH | ** | 0.0061 |
| PBS vs. VEH\|SIV | ** | 0.0035 | PBS vs. VEH\|SIV | ** | 0.0038 |
| PBS vs. THC\|SIV | ns | 0.3623 | PBS vs. THC\|SIV | ns | 0.3955 |
| DEX vs. TNFα | ns | 0.0605 | DEX vs. TNFα | * | 0.0271 |
| DEX vs. VEH | *** | 0.0004 | DEX vs. VEH | ** | 0.0036 |
| DEX vs. VEH\|SIV | **** | <0.0001 | DEX vs. VEH\|SIV | ** | 0.0030 |
| DEX vs. THC\|SIV | * | 0.0251 | DEX vs. THC\|SIV | * | 0.0424 |
| TNFα vs. VEH | ns | 0.3233 | TNFα vs. VEH | ns | 0.1021 |
| TNFα vs. VEH\|SIV | ns | 0.6995 | TNFα vs. VEH\|SIV | ns | 0.3892 |
| TNFα vs. THC\|SIV | ns | 0.0951 | TNFα vs. THC\|SIV | * | 0.0420 |
| VEH vs. VEH\|SIV | * | 0.0271 | VEH vs. VEH\|SIV | ns | 0.0666 |
| VEH vs. THC\|SIV | ** | 0.0090 | VEH vs. THC\|SIV | * | 0.0178 |
| VEH\|SIV vs. THC\|SIV | *** | 0.0002 | VEH\|SIV vs. THC\|SIV | * | 0.0102 |

**Figure 4.**
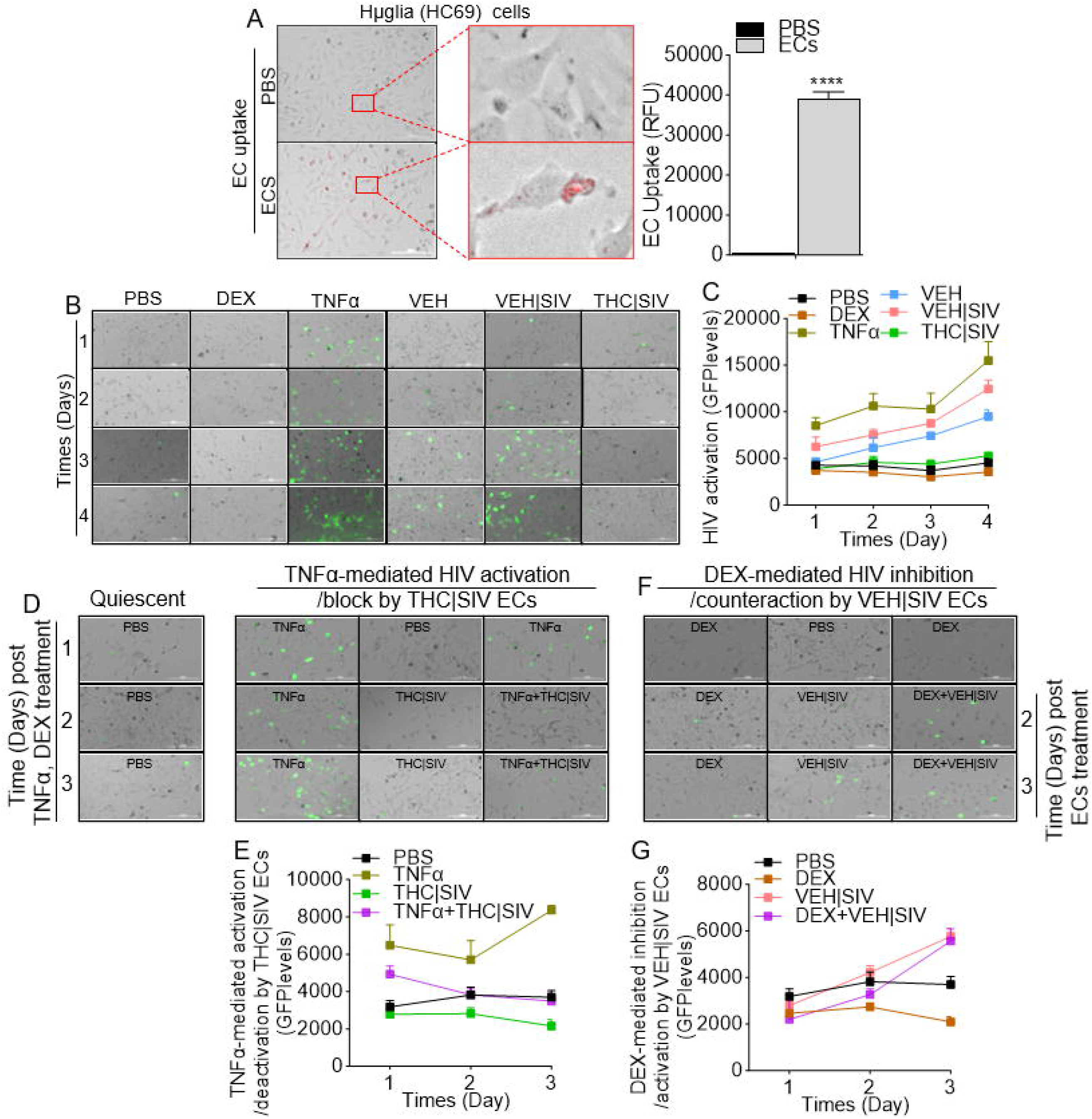
Effect of basal ganglia derived ECs on HIV reactivation in microglia (hµglia, HC69) cells. **A**) Representative microscopic images of internalized DIR-labeled ECs by huglia cells (left) and the level of internalized ECs (bars, right). **B**) Representative microscopic images of huglia-GFP cells showing HIV reactivation (GFP expression, green) at different times after treatment with VEH, VEH|SIV, THC|SIV ECs and other compounds (PBS, DEX, TNFα). **C**) The levels of HIV reactivation (GFP expression). **D**) Representative microscopic images of hμglia-GFP cells showing THC|SIV suppressing TNFα-mediated HIV reactivation. **E**) The levels of HIV reactivation (GFP expression) by TNFα and suppression by THC|SIV ECs. **F**) Representative microscopic images of hμglia-GFP cells showing VEH|SIV activating DEX-mediated HIV reactivation. **G**) The levels of HIV suppression (GFP expression) by DEX and reactivation by VEH|SIV ECs. All experiments were repeated three times. Statistical differences were assessed by ordinary 2-way ANOVA with Tukey’s multiple comparison test as presented on Tables 4 – 6. ns = non-significant.

### THC|SIV ECs restrict the reactivation of latent HIV in microglia

Here, we assessed the effects of ECs on TNFα-mediated HIV activation and DEX-mediated HIV inhibition in microglia. Latently infected hµglia cells were treated with PBS, THC|SIV, VEH|SIV, or stimulated with TNFα (10 ng/mL), Dex (3 mM/mL). Cells were cultured for 1 day. On day 2, THC|SIV ECs were added to TNFα-treated cells while VEH|SIV ECs were added to DEX-treated cells. Reactivation of latent proviruses was evaluated at different time points after treatment (days 1, 2, 3) by measuring GFP expression. Strong inhibition of HIV proviral reactivation in TNFα-stimulated cells was observed in cells treated with THC|SIV treatment (**Fig. 4D, 4E, Table 5**). In contrast, strong HIV proviral reactivation in DEX-suppressed cells was observed in cells treated with VEH|SIV ECs (**Fig. 4F, 4G, Table 6**). The levels of GFP expression in TNFα+THC|SIV is similar to THC|SIV treated cells and significantly lower than TNFα treated cells (**Fig. 4D, 4E, Table 5**), indicating that THC|SIV ECs suppressed the ability of TNFα to reactivate HIV. In parallel, the levels of GFP expression in DEX+VEH|SIV is similar to VEH|SIV treated cells and significantly higher than DEX treated cells (**Fig. 4F, 4G, Table 6**), indicating that VEH|SIV ECs countered the ability of DEX to suppress latent HIV activation.

**Table 5.**
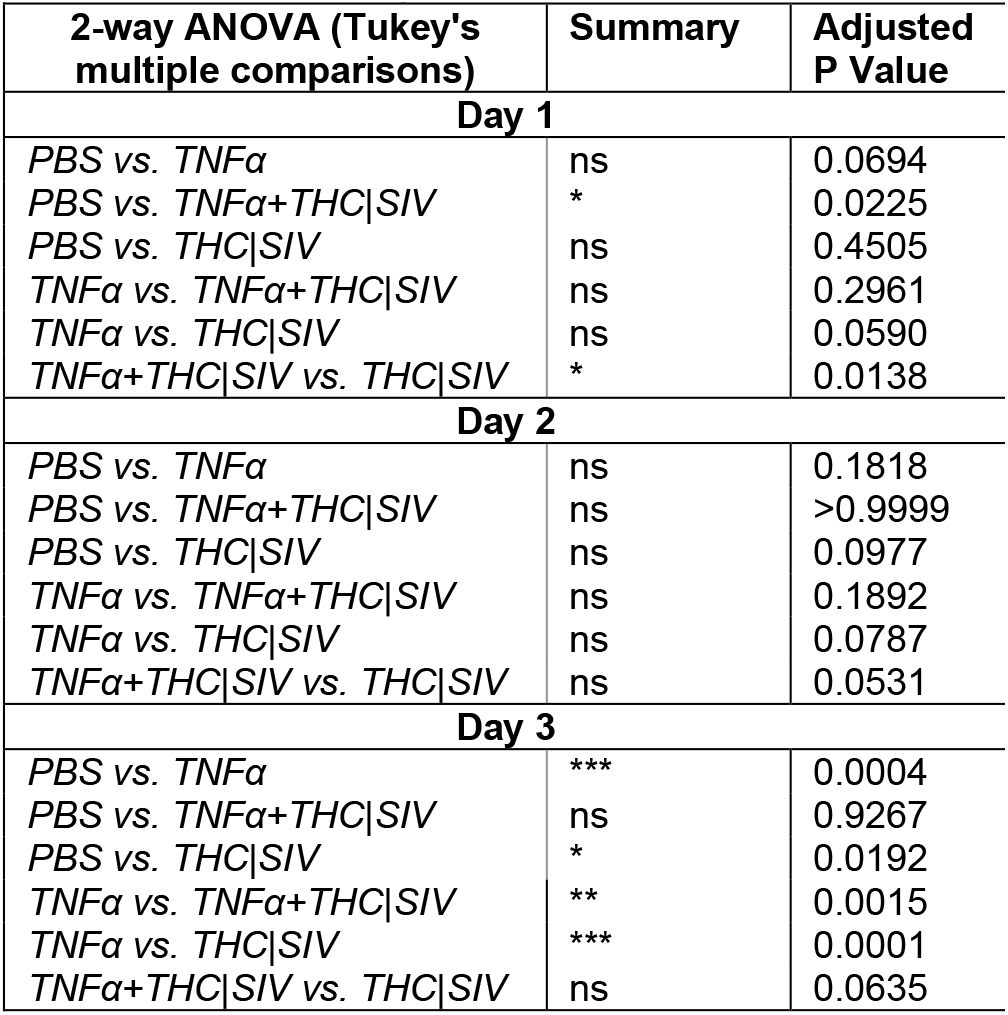
Statistics for TNFα-mediated activation/deactivation by THC|SIV ECs (GFP levels)

**Table 6.**
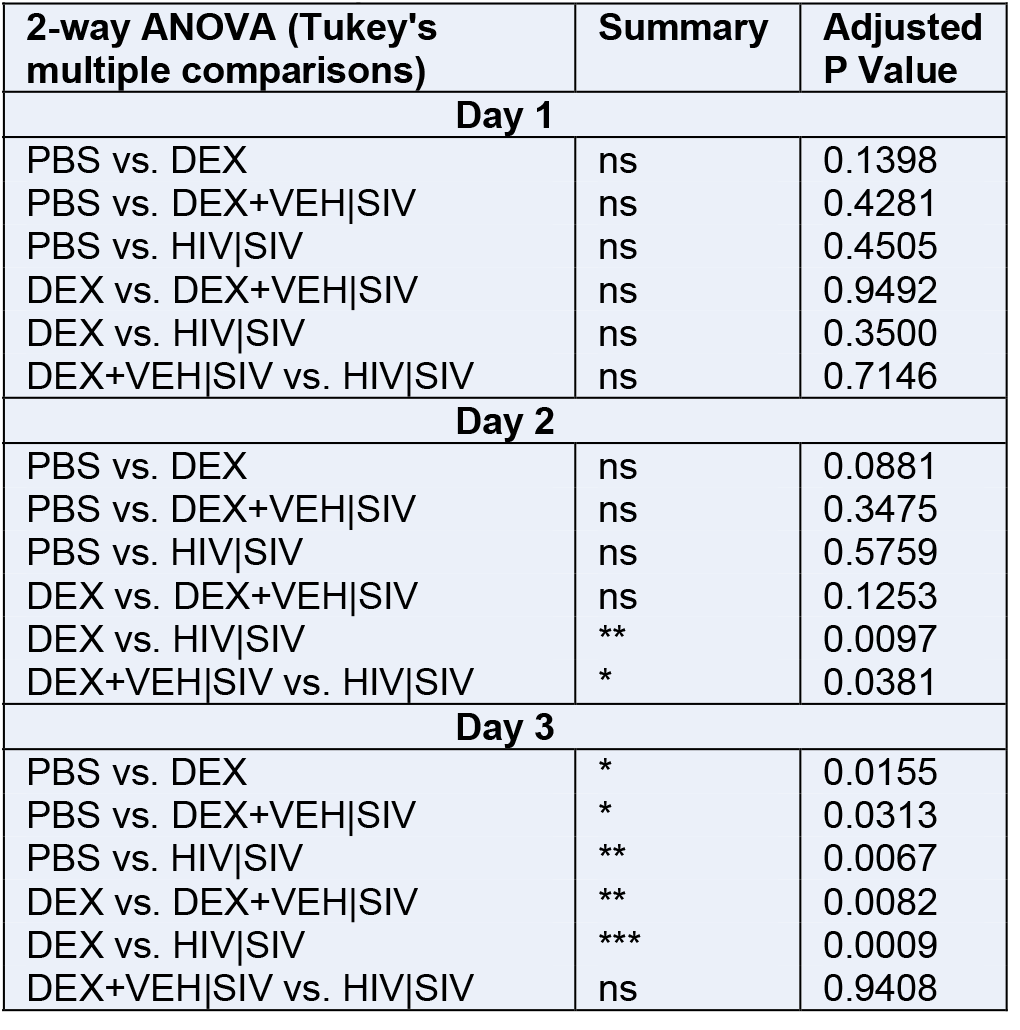
Statistics for DEX-mediated inhibition/activation by VEH|SIV ECs (GFP levels)

### ECs reprogram transcriptome of HIV latently infected monocytes

We mapped U1 RNA-seq reads to RefSeq-annotated human gene loci, quantified reads as transcripts per million mapped reads (TPM) and genes are considered expressed if TPM >1 in all three samples within a treatment group. Data filtered with p-value (unpaired t-test) <= 1.0 and FC >= 0.0, resulted in 14440 mRNAs (**Fig. 5A, Supplemental Table 1**). Differentially expressed genes (DEGs) identified using p-value (unpaired t-test) <0.05, p-adj (FDR <0.05) were 898, 507, 943 for VEH, VEH|SIV, and THC|SIV respectively (**Supplemental Table 2**). 3-way Venn overlap analysis identified EC-altered genes that are either unique or common to all 3 treatments (**Fig. 5B**, **Supplemental Table 3**). The expression levels of the common genes are shown on the heatmap (**Fig. 5C**). Of the 13 common genes, VEH|SIV ECs upregulated 8 and down regulated 5 genes while THC|SIV ECs had opposite effect. Of interest was the upregulation by VEH|SIV ECs and downregulation by THC|SIV ECs of LLGL2 and PTPRN2. LLGL2 forms a complex with GPSM2/LGN, PRKCI/aPKC and PARD6B/Par-6 to ensure the correct organization and orientation of bipolar spindles for normal cell division. PTPRN2 is a receptor-type tyrosine-protein phosphatase N2 required for normal accumulation of secretory vesicles in hippocampus, normal accumulation of neurotransmitters (norepinephrine, dopamine, serotonin) in the brain. Also of interest is the differential effect of ECs on PLXNB1, which was downregulated by VEH|SIV ECs and upregulated by THC|SIV ECs. PLXNB1 is a receptor for SEMA4D, which plays a role in GABAergic synapse development, mediates SEMA4A- and SEMA4D-dependent inhibitory synapse development, and plays a role in RHOA activation and subsequent changes of the actin cytoskeleton, axon guidance, invasive growth and cell migration. Similarly, AURKB was downregulated by VEH|SIV ECs and upregulated by THC|SIV ECs. Interestingly, AURKB is a key cell cycle regulatory kinase known to inhibit HIV cell-to-cell transmission [27], similar to the effect of THC|SIV ECs that inhibits cell-to-cell transmission (**Figs. 3Q, R**). Overrepresentation analysis of the common genes identified important gene ontology (GO) biological processes associated with the 13 common genes in **Fig. 5C**, where 11 genes were annotated to the GO biological process and used for the enrichment analysis (**Fig. 5D**).

**Figure 5.**
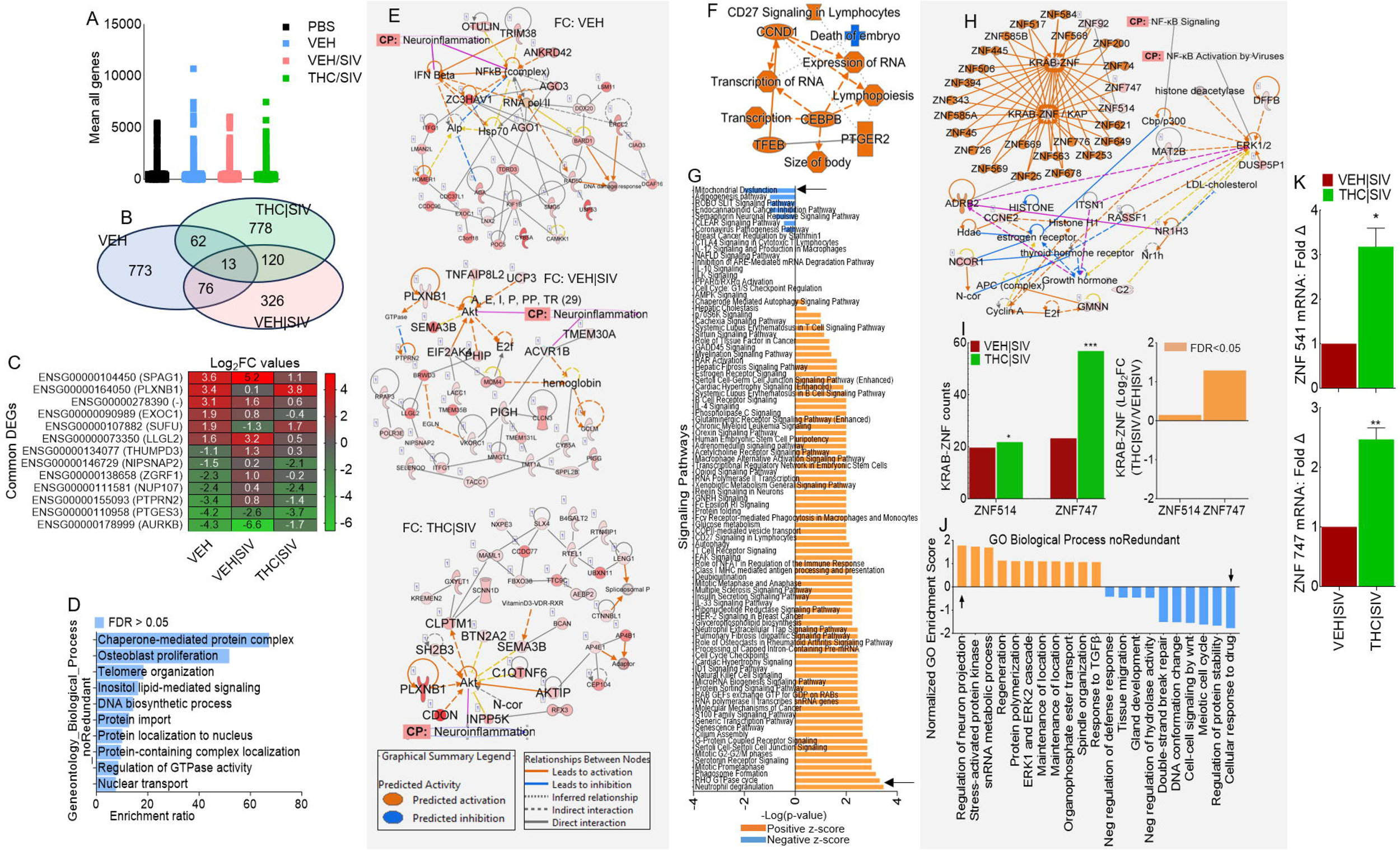
Global transcriptome analysis of HIV latently infected U1 cells treated with basal ganglia derived ECs. **A**) Scatter plot of all transcripts from biological triplicates with transcripts per million mapped reads (TPM) >1, p-value (unpaired t-test) <= 1.0, and fold-change >= 0.0 are displayed. **B**) 3-way Venn diagram of DEGs meeting the following criteria: i) pooled-adjusted p-value (FDR) < 0.05 and ii) (log_2_ fold change >1 or <-1). **C**) Heatmap of the genes corresponding to the 13 common DEGs identified by the 3-way Venn diagram. **D**) Overrepresentation analysis of gene ontology biological processes of the 13 common DEG identified by the 3-way Venn diagram. The y-axis represents and the x-axis represents enrichment ratios. **E**) IPA network diagrams representing the regulatory effects with top consistency scores found associations with neuroinflammation signaling, with the top network representing FC: VEH/PBS, middle network representing FC: VEH|SIV/PBS, and bottom network representing FC: THC|SIV/PBS. FC = Fold Change. **F**) Gene to function analysis of THC|SIV/VEH|SIV DEGs. **G**) Histogram of the 92 canonical pathways regulated by treatment with THC|SIV ECs. The left y-axis represents the canonical pathways, and the x-axis represents -log (P-value) the canonical pathways. Black arrows point to the most upregulated (neutrophil degranulation) and downregulated (mitochondrial dysfunction) pathways. Each bar indicates the value of -log_10_ (P-value). **H**) IPA network diagram representing top consistency scores with links to NF-kB activation by viruses and signaling, as well as KRAB-ZNF (Krüppel-associated box domain zinc finger) gene cluster. **I**) Differences in ZNF514 and ZNF747 represented as counts (left) and log_2_ fold change (right). **J**) Gene ontology gene set enrichment analysis (GSEA). The left y-axis represents normalized GO enrichment scores, and the x-axis represents GO biological process (noRedundant). **K**) DEGs validated with RT-qPCR analysis. Statistical differences were assessed by ordinary one-way ANOVA with Tukey’s correction and by Binary Student’s t tests (Welch’s correction). **** p < 0.001, *** p < 0.005, ** p < 0.01, * p < 0.05, and ns = non-significant.

### Gene-to-function analysis of the transcriptome of EC-treated cells predicts links to neuroinflammation

IPA pathway and network analyses of genes in **Figure 5B** showed that ECs regulate various canonical pathways, biological, cellular, diseases, and molecular functions (**Table 7**). The different ECs are predicted to regulate biological processes with neurological disease as the first of the top 5 diseases for VEH|SIV treated cells and the last of the top 5 diseases for THC|SIV treated cells (**Table 7**). Treatment with VEH ECs is linked to 11 networks that have 29 focused molecules with activated IFNβ and NF-KB complex as hub networks driving neuroinflammation signaling (**Fig. 5E, top, Supplemental Fig. 2**). VEH|SIV treated cells had 9 networks with 30 focused molecules with activated Akt and ACVR1B as hub networks driving neuroinflammation signaling (**Fig. 5E, middle, Supplemental Fig. 3**). On the other hand, THC|SIV treated cells with 20 networks with 30 focused molecules and activated Akt as a hub network driving neuroinflammation signaling (**Fig. 5E, bottom, Supplemental Fig. 4**). Detailed IPA analysis revealed that VEH ECs may regulate neuroinflammation by downregulating NF-KB complex leading to inhibition of TGFβ, neuronal, and microglia survival, and increased accumulation of nitric oxide, while activating IFNβ signaling, which leads to increased myelin debris clearance and inhibition of activated MHC Class II (**Supplemental Fig. 2**). Although VEH|SIV and THC|SIV ECs regulate neuroinflammation via Akt, the Akt-mediated regulation of neuroinflammation function through different executioner molecules. VEH|SIV ECs may regulate neuroinflammation via Akt interaction with glycogen synthase kinase 3 beta (GSK3B) mediated inhibition of NF-kβ activation that leads to inhibition of TGFβ, inhibition of neuronal and microglia survival, and increased accumulation of nitric oxide, inhibition of activated MHC Class II, as well as GSK3B mediated blockade of AP1 (**Supplemental Fig. 3**). Unlike VEH|SIV ECs, THC|SIV ECs mediate neuroinflammation via Akt interaction with GSK3B mediated activation of NF-kβ signaling that leads to activation of TGFβ, increased neuronal and microglia survival, and decreased accumulation of nitric oxide and inhibition of MHC Class II, as well as GSK3B mediated blockade of activated AP1 (**Supplemental Fig. 4**). These findings are significant because Akt-induced NF-kB via GSK3B may promote or impair survival of latently infected cells.

**Table 7.** Three Biological Processes induced by different EC treatment.

| Group | Top 5 Canonical Pathways |  |
| --- | --- | --- |
|  | Name | p-value range |
| VEH | RNA Pol III Transcription | 1.25E-04 |
|  | Assembly of RNA Pol III Complex | 6.19E-03 |
|  | mRNA Capping | 1.19E-02 |
|  | Nucleotide Excision Repair Pathway | 1.44E-02 |
|  | Nucleosome assembly | 2.01E-02 |
| VEH\|SIV | Metabolism of vitamin K | 5.29E-03 |
|  | PI Metabolism | 9.47E-03 |
|  | Xanthine and Xanthosine Salvage | 1.06E-02 |
|  | Cell Cycle Checkpoints | 1.50E-02 |
|  | Guanine and Guanosine Salvage | 1.58E-02 |
| THC\|SIV | PI Metabolism | 2.49E-03 |
|  | Mitotic Roles of Polo-Like Kinase | 7.25E-03 |
|  | Cilium Assembly | 8.51E-03 |
|  | NAD Phosphorylation and Dephosphorylation | 9.32E-03 |
|  | Gamma carboxylation, hypusine formation and arylsulfatase activation | 1.14E-02 |
| Top 5 Diseases and Biological Functions |  |  |
| Group | Name | p-value range |
| VEH | Connective Tissue Disorders | 4.14E-04 - 4.14E-04 |
|  | Developmental Disorder | 4.14E-04 - 4.14E-04 |
|  | Hereditary Disorder | 2.21E-02 - 4.14E-04 |
|  | Organismal Injury and Abnormalities | 4.80E-02 - 4.14E-04 |
|  | Skeletal and Muscular Disorders | 2.13E-02 - 4.14E-04 |
| VEH\|SIV | Neurological Disease | 4.92E-02 - 5.77E-07 |
|  | Organismal Injury and Abnormalities | 4.92E-02 - 5.77E-07 |
|  | Cancer | 4.92E-02 - 3.41E-06 |
|  | Dermatological Diseases and Conditions | 4.67E-02 - 5.75E-04 |
|  | Gastrointestinal Disease | 4.67E-02 - 9.10E-04 |
| THC\|SIV | Cancer | 4.49E-02 - 7.36E-17 |
|  | Organismal Injury and Abnormalities | 4.49E-02 - 7.36E-17 |
|  | Gastrointestinal Disease | 4.27E-02 - 1.10E-13 |
|  | Endocrine System Disorders | 4.42E-02 - 4.83E-10 |
|  | Neurological Disease | 4.49E-02 - 1.14E-07 |
| Top 5 Molecular and Cellular Functions |  |  |
| VEH | **Name** | **p-value range** |
|  | Cell Death and Survival | 3.54E-02 - 4.14E-04 |
|  | Cellular Assembly & Organization | 2.74E-02 - 4.14E-04 |
|  | DNA Replication, Recombination, Repair | 2.13E-02 - 4.14E-04 |
|  | Small Molecule Biochemistry | 2.13E-02 - 8.27E-04 |
|  | Cell Cycle | 2.74E-02 - 1.24E-03 |
| VEH\|SIV | Cell Death and Survival | 4.25E-02 - 5.77E-07 |
|  | Cell Morphology | 4.67E-02 - 5.77E-07 |
|  | Cellular Compromise | 3.13E-02 - 5.77E-07 |
|  | Cellular Assembly and Organization | 4.67E-02 - 1.77E-03 |
|  | Cellular Function and Maintenance | 4.67E-02 - 1.77E-03 |
| THC\|SIV | Cell Morphology | 4.49E-02 - 1.30E-04 |
|  | Cell Death and Survival | 4.49E-02 - 6.51E-04 |
|  | Cellular Assembly and Organization | 4.49E-02 - 1.10E-03 |
|  | Cellular Function and Maintenance | 4.49E-02 - 1.10E-03 |
|  | Carbohydrate Metabolism | 4.49E-02 - 1.89E-03 |

### THC|SIV ECs regulate expression of KRAB-ZNF gene family

We identified 745 DEGs that are significantly modified by THC|SIV (**Supplemental Table 2**). Gene-to-function analysis identified 9 upregulated and 1 downregulated biological process (**Fig. 5F**). Neutrophil degranulation and mitochondrial dysfunction are the most upregulated and downregulated pathways, respectively (**Fig. 5G, black arrows**). 19 IPA biological networks were identified. Merging of networks 8 and 17 linked to neurological disease identified KRAB-ZNF cluster containing 24 family members and linked to ERK1/2 activation and NF-kB activation via ADRB2 (**Fig. 5H**), a regulator of the neuroimmune response [28]. Of the 24 ZNF family members, ZNF514 and ZNF747 are present in our data set, and both were upregulated by THC|SIV ECs (**Fig. 5I**). GO gene set enrichment analysis (GSEA) of KRAB-ZNF cluster predicted that the THC|SIV treatment is associated with GO terms that include increased regulation of neuron projection development and decreased cellular response to drugs among others (**Fig. 5J**). ZNF514 and ZNF747 expression (**Fig. 5K**) were validated with RT-qPCR using primer sequences in **Table 2**.

### THC|SIV ECs augment secretion of TH2 and suppression of inflammasome cytokines

Secretome analysis of supernatants from U1 cells treated with PBS or ECs identified 105 secreted proteins (**Fig. 6A**). 34, 91, 68, and 81 proteins were significantly altered in VEH, VEH|SIV, THC|SIV, and THC|SIV/VEH|SIV (**Fig. 6B**). 4-way Venn overlap analysis identified proteins that are common amongst groups or those that are unique to each group (**Fig. 6C**). The fold change differences amongst all the common proteins are displayed as bar graphs (**Fig. 6D**). Network analyses of significantly different proteins identified the main hub proteins of each network (**Fig. 6E**). The levels (**Fig. 6F**) and PPI network of the top 5 upregulated (**Fig. 6G, left**) and top 5 downregulated (**Fig. 6G, right**) proteins were determined. Upregulated proteins are in 3 clusters with the Th2 cytokines IL-5, IL-10, IL-24 in a cluster linked to GO biological process related to regulation of signaling pathway via JAK-STAT. Downregulated proteins are in 3 clusters with IL-13, IL-18, LEP in a cluster linked to GO biological process related to regulation of natural killer cell proliferation and inflammatory bowel disease in KEGG pathway. Secretion of IL-5, IL-10, IL-24, and IL-12, but not sST2 were confirmed in supernatant from hµglia cells using ELISA (**Fig. 6H**). In parallel, IL-18, IL-23, Leptin, EGF, but not IL-13 was validated by ELISA in supernatant from hµglia cells (**Fig. 6I**).

**Figure 6.**
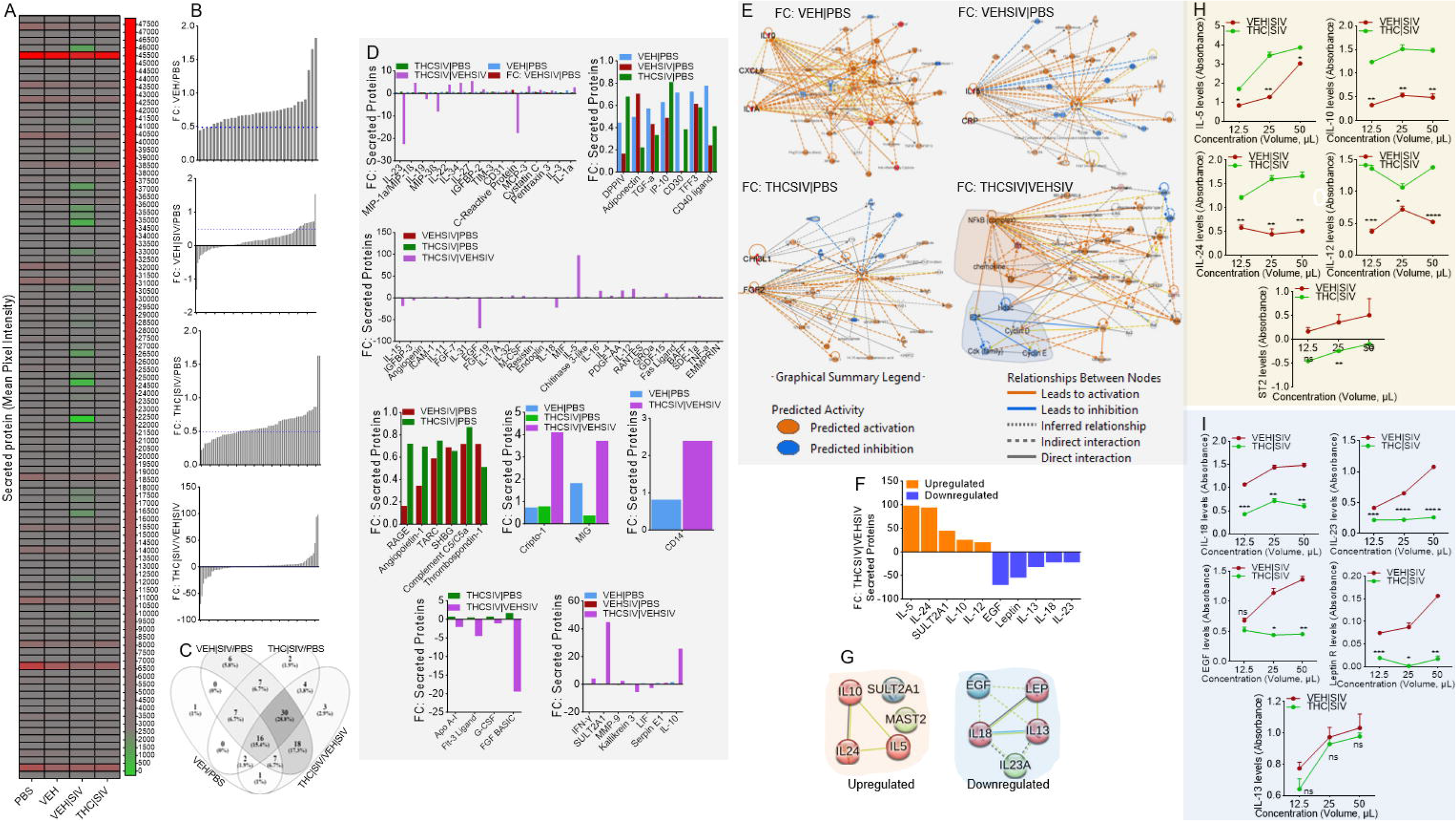
Altered secretion of proteins by HIV latently infected U1 cells treated with basal ganglia derived ECs. **A**) Heatmap of Mean Pixel Intensity of identified secreted proteins. **B**) Fold change (FC) of secreted proteins between groups. From top to bottom are plots for VEH/PBS (34 proteins), VEH|SIV/PBS (91 proteins), THC|SIV/PBS (68 proteins), THC|SIV/VEH|SIV (81 proteins). The y-axis represents DEG fold change (FC), while the x-axis represents altered DEGs. **C**) 4-way Venn diagram-based overlap analysis of all secreted proteins showing the number of overlapping and unique proteins. **D**) The plot of the fold change differences amongst all the common proteins are plotted as bar graphs. The y-axis represents Fold Change, and the x-axis represents enriched proteins. **E**) IPA network diagrams representing the regulatory networks with top consistency scores found associations with different molecules. **F**) Levels of top 10 differentially secreted proteins between THC|SIV/VEH|SIV. The y-axis represents Fold Change, and the x-axis represents differential proteins. **G**) STRING interactome analysis of upregulated (left) and downregulated (right) proteins. The blue, green, red nodes represent different clusters. The line thickness indicates the strengths of the interactions. **H-I**) The preferential secretion of the proteins in panel “F” was validated by ELISA analysis. Four (IL-5, IL-10, IL-24, IL-12) out of the five upregulated proteins were validated (**H**). Three (IL-18, EGF, Leptin) out of the five downregulated proteins were validated (**I**).

## Discussion

In this study, we showed that ECs are previously unknown endogenous latency regulating agents. VEH|SIV BG ECs potently induced durable activation of latent HIV in three different cell line models (J-Lat GFP/J-Lat Tat-GFP T cells, U1 monocytes, HC69 microglia) of HIV latency as evidenced by transactivation of HIV LTR promoter, expression of cell-associated HIV mRNA, and release of infectious virions. In contrast, THC|SIV ECs either did not or minimally activated latent HIV in the three cell lines (**Figs. 1-4**). Virions produced by EC-treated cells induced syncytia formation both in short- and long-term assays and were infectious as they mediated cell-free and cell-to-cell viral spread in the following order VEH|SIV > VEH > THC|SIV = PBS. The induction of syncytium or lack thereof by VEH|SIV or THC|SIV, respectively are significant because syncytium is a form of cell fusion that likely forms due to excess karyogamy -nuclear fusion within the syncytium [29], where giant cells are formed by infected cells fusing with neighboring uninfected cells, as a means of disseminating the virus [30, 31]. The inhibitory effects of THC|SIV ECs suggest that THC|SIV ECs do not have proinflammatory cargo to activate latently infected cells or that the THC|SIV ECs could induce anti-inflammatory responses. may drive HIV into ‘super latency’. THC|SIV ECs-mediated promotion of latency is durable because it blocked HIV transcription up to 16 days with or without additional treatment (**Fig. 3**) and failed to reactivate latent HIV despite treatment with potent reactivators like TNFα (**Figs. 4D-G**).

ECs mediate specific transcriptional profiles and gene networks that regulate neuroinflammation. In the presence of VEH and VEH|SIV ECs, Akt activation may result in loss of NF-kB activation via GSK3B while THC|SIV ECs may result in stimulation of NF-kB by Akt (**Supplemental Figs. 2-4**). These results demonstrate two separate functions of the Akt complex in NF-kB activation in HIV latently infected cells. Furthermore, THC|SIV EC-mediated activation of KRAB-ZNF gene cluster is remarkable but their contribution to HIV pathogenesis remains unclear. Activation of KRAB-ZNF cluster (**Fig. 5H**) is interesting because KRAB-ZNF may have pleiotropic effects on transcriptional regulation of its target genes [32] since they bind target promoters via specific DNA recognition sequences and regulate transcription by RNA pol II. In HIV infected cells, intact proviruses preferentially integrate within KRAB–ZNF genes and ZNF genes carrying clonal intact integrations are down-regulated upon cellular activation [33]. ZNF304 silences HIV gene transcription via recruitment to the viral promoter of heterochromatin-inducing methyltransferases [34]. KRAB-ZNF proteins function as potent transcriptional repressors and thus, in addition to the local chromatin environment, KRAB-ZNF proteins may be critical regulators of viral latency. Whether or not ZNF514 and ZNF747 genes regulate HIV latency remains to be determined.

Of note, the secretome profile of cells treated with THC|SIV ECs clearly distinguished functions of the ECs in suppressing reactivation of latent proviruses. In the setting of THC|SIV ECs, gene-to-function analyses revealed activated NF-kB complex, which is a hub for IFN, TLR, and chemokine activation that may contribute to the control of virus replication. Causal network analysis of i) NF-kB activation by viruses via ERK1/2, and ii) NF-kB signaling via Cbp/p300, may mediate gene expression by regulating chromatin structure at the gene promoter through their intrinsic histone acetyltransferase (HAT) activity. Such activities may promote or block the recruitment of basal transcriptional machinery including RNA pol II to the HIV promoter. Interestingly, THC|SIV ECs preferentially increased the levels of Th2 cytokines IL-5, IL-10, and IL-24, but decreased the levels of proinflammatory molecules IL-18, IL-23, Leptin, and EGF. Pro-inflammatory properties of leptin are similar to those of the acute phase reactants and upregulates the secretion of inflammatory cytokines [35]. The proinflammatory cytokine IL-18, that is mainly secreted by myeloid cells plays an important role in host response to infection by viruses and intracellular pathogens [36]. IL-18 is elevated in the serum of PLWH [37], increased during ART failure but decreased in virally suppressed individuals [38]. THC|SIV ECs-mediated suppression of IL-18 secretion, inhibition of HIV transcription, and production of infectious virions suggest that IL-18 may regulate HIV persistence by enhancing reactivation of latent proviruses and/or increasing HIV replication, as suggested for HIV infected monocytes and T cell lines [39-41]. The current findings support our previously published studies showing that treatment of SIV-infected macaques with THC results in suppressed inflammation and secretion of extracellular vesicles (EVs) with anti-inflammatory functions [9-11, 15, 16, 42].

The ability of latently HIV infected cells to respond to stimulation with VEH ECs but more potently by VEH|SIV ECs suggest that these endogenous ECs can reverse HIV latency in vitro. In the absence of ART or failed treatment, these ECs may provide a persistent source of viremia in vivo. On the other hand, the Th2-biased secretome signature (increased secretion of IL-5, IL-10, IL-24) by THC|SIV EC treated monocytes and microglia in addition to the inhibition of HIV transcription support the notion that THC|SIV ECs inhibit reactivation of latent HIV and possibly suppress inflammation. While there is no consensus on the role of IL-10 in HIV pathogenesis [43], increased levels of IL-10 was shown to inhibit HIV replication *in vitro* [44] and elevated plasma levels of IL-10 was associated with the control of viral replication in pregnant women [45]. Although increased levels of IL-10 in PLWH may be seen as deleterious due to the potential of IL-10 to decrease the production of Th1 cytokines and skew host response towards Th2 [46], there is broad unanimity in the HIV field that progression of HIV and non-HIV diseases is closely linked to persistent immune activation.

The recognition that EVs and ECs (**Figs. 1-6**) isolated from THC treated SIV-infected RMs have the ability to suppress virus-induced inflammation and inhibit virus production from latently-infected cells emphasizes the importance of understanding the antiviral roles of EVs and ECs in the setting of HIV cure research. To the best of our knowledge, this current study represents an initial and most comprehensive investigation focusing on the potential role of ECs on HIV persistence and the impact of THC in regulating viral persistence via ECs.

Future studies are required to identify the EC cargo that mediates these effects to better understand the cellular factors that reactivate latently HIV/SIV infected cells in the brain.

## Methods

Detailed methods are presented in supplemental materials.

### Purification and characterization of BG EC

ECs used in this study were previously characterized [11].

### Cells lines

TZM-GFP [47]; HIV-1 infected U937 Cells (U1) [48]; J-Lat GFP and J-Lat Tat-GFP cells [49, 50] were obtained through the NIH HIV Reagent Program. Hμglia (HC69) and CEMx174 cells were kind gifts from Drs. Jonathan Karn and Mahesh Mohan respectively.

### EC internalization

PBS or ECs were stained with 5 µM SYTO™ RNASelect™ or DiR’; DiIC18(7) (1,1’-Dioctadecyl-3,3,3’,3’-Tetramethylindotricarbocyanine Iodide) [51-54]. Images of the cells were taken at different times, processed (Gen5), and plotted (GraphPad Prism 10.1) [16, 24, 25].

### RT-qPCR

Five µg total RNA was used for cDNA synthesis and real-time PCR (**Table 2)** [55].

### Western Blot

A total of 50 µg of protein extracts from U1 cells treated with PBS or ECs for 4 days were subjected to 4–20% SDS-PAGE and western blot with relevant antibodies [23, 25, 56]. Blots were processed and images were captured using LI-COR, and band intensity measured using ImageJ [23, 25, 56].

### Viability Assay

10,000 cells/well were seeded in 96-well plates and treated with ECs (50 µg/mL) or equivalent volume of 0.1× PBS for 04 days at 37 °C. On day 04 of treatment, cells were counted, and the number of live cells determined using the Luna-II automated cell counter and validated using the trypan blue exclusion assay.

### RNA-Seq

100-300 µg total RNA were isolated, and RNA quality was assessed by RNA Tapestation and quantified by Qubit 2.0 RNA HS assay [55]. Prior to first strand synthesis, samples were randomly primed (5′ d(N6) 3′ [N=A, C,G,T]) and fragmented based on manufacturer’s recommendations. The first strand is synthesized with the Protoscript II Reverse Transcriptase with a longer extension period, approximately 40 minutes at 42°C. All remaining steps for library construction were used according to the NEBNext® Ultra™ II Directional RNA Library Prep Kit for Illumina®. Final libraries quantity was assessed by Qubit 2.0 and quality was assessed by TapeStation D1000 ScreenTape. Equimolar pooling of libraries was sequenced on Illumina® Novaseq platform with a read length configuration of 150 PE for 40M PE reads/sample (20M in each direction).

### Bioinformatics

FastQC was applied to check the quality of raw reads. Trimmomatic was applied to cut adaptors and trim low-quality bases with default setting. STAR Aligner was used to align the reads. The package of Picard tools was applied to mark duplicates of mapping. StringTie was used to assemble the RNA-Seq alignments into potential transcripts. FeatureCounts or HTSeq was used to count mapped reads for genomic features such as genes, exons, promoter, gene bodies, genomic bins, and chromosomal locations. Raw TPMs [57, 58] are provided in **Supplemental Table 1**.

### Secretome analysis

100,000 U1 cells were seeded in a 12 well plate with 500 uL of RPMI media. Cells were treated with ECs (50 ug) or 0.1X PBS for 04 days. On day four, cells were collected, processed, pellets saved for transcriptome analysis. 1 mL of clarified supernatants from each group was used to perform cytokine array analysis with Proteome Profiler Human XL Cytokine Array Kit per manufacturer’s protocol. Target protein expression as Dot blot were quantified using Empiria Studio and protein expression presented as pixel intensity.

### Syncytia Assay

5000 CEMx174 cells/well were seeded in 48 well plate with 250 µL of RPMI media. After two hours, 250 µL of clarified supernatants from PBS or ECs treated U1 cells were added to the cells. Cells were cultured at 37 °C and syncytia were visualized on days 1, 2, and 4 at 10X using Lionheart FX microscope.

### Cell to cell HIV transfer

Cell-to-cell infection of TZM-GFP cells was performed with 1000 PKHRed-labeled CEMx174 cells cultured in the presence of 250 µL of clarified supernatants from PBS or ECs treated U1 cells. After four days, the PKHRed-labeled CEMx174 were overlayed atop 1000 TZM-GFP cells and the co-culture incubated for 2 days. The activation of GFP expression in TZM-GFP cells is a sign of virus transfer from CEMx174.

### Validation of transcriptome and secretome data

HC69 Huglia cells were used for data validation with RT-qPCR [55] using total RNA cells ELISA [59] using supernatants.

### Flow Cytometry

Cells of interest were processed, resuspended in MACSQuant running buffer, and GFP expression data acquired using BD Flow cytometer and analyzed using FlowJo™ as previously described [59].

### Statistical analysis

Significance cutoff was set to fold change (FC) >1.5 or <-1.5 and p-value <0.05. Statistical differences were assessed by one-or two-way ANOVA with Šídák’s or Tukey’s multiple comparisons test, or Binary Student’s t tests (Welch’s correction) using GraphPad Prism 10. Details of specific statistics are on figure legends, as well as in **Tables 2 to 4**.

## Supporting information

Suplemental Table 1 List of genes with Pvalues 1

## Declarations

### Consent for publication

All authors read and approved the publication of this manuscript.

### Data availability

RNA-Seq and Secretome datasets are included within the article and its additional files.

### Author contributions

CMO and MM conceptualized the study. WN, and LSP assisted with nonhuman primate sample collection, tissue processing and performed the experiments. WN, BCO, MM, and CMO conducted data analyses. WN, BCO, MM, and CMO wrote the original draft of the paper, provided text, reviewed and edited the manuscript.

## Acknowledgments

This work was supported, in part, by the New York Medical College Imaging Core, Iowa Hybridoma Bank, University of Iowa, Texas Biomedical Research Institute.

## Figure legends

**Supplemental Figure 1:** A) Description of cell lines for model systems of post-integration latency. B-D) Pictorial description of cell types, treatments, and length of treatment and culturing.

**Supplemental Figure 2:** Ingenuity Pathway Analysis (IPA) neuroinflammation signaling pathway induced by ECs isolated from control VEH rhesus macaques. Shown are the principal interaction networks in cells treated with VEH ECs.

**Supplemental Figure 3:** Ingenuity Pathway Analysis (IPA) neuroinflammation signaling pathway induced by ECs isolated from SIV-infected (VEH|SIV) rhesus macaques. Shown are the principal interaction networks in cells treated with VEH|SIV ECs.

**Supplemental Figure 4:** Ingenuity Pathway Analysis (IPA) neuroinflammation signaling pathway induced by ECs isolated from THC-treated SIV-infected (THC|SIV) rhesus macaques. Shown are the principal interaction networks in cells treated with THC|SIV ECs.

