## Supplemental Figures for "Extracellular condensates (ECs) are endogenous modulators of HIV transcription and latency reactivation"

### Supplemental Figure 1

A

| Cell line | Replication competent virus | Site of mutation | Proviral copies/cell | Reference |
|---|---|---|---|---|
| J-Lat GFP | No | LTR-GFP | 1 | Jordan et al., 2001; Jordan et al., 2003. |
| J-Lat Tat-GFP | No | LTR-Tat-GFP | 1 | Jordan et al., 2001; Jordan et al., 2003. |
| U1 | Yes | Tat | 2 | Folks et al., 1989 |
| Hμglia (HC69) | Yes | Tat | 1 | Alvarez-Carbonell et al., 2017 |

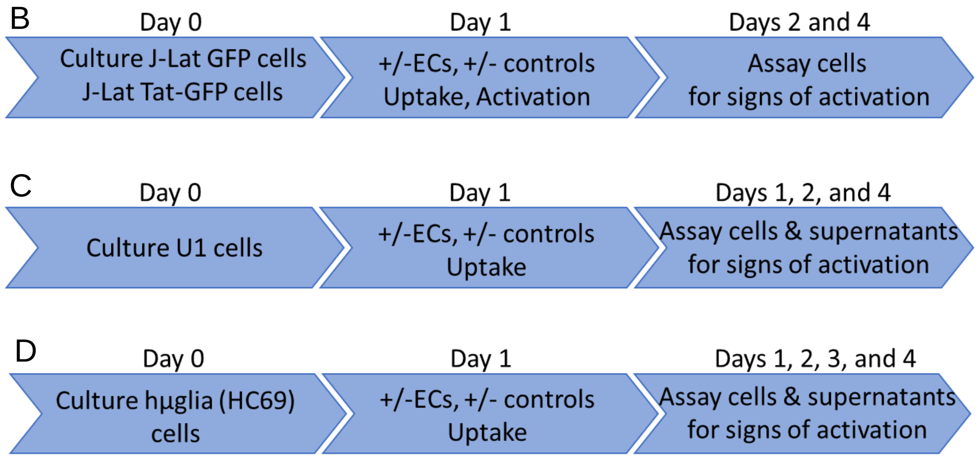

### Supplemental Figure 2

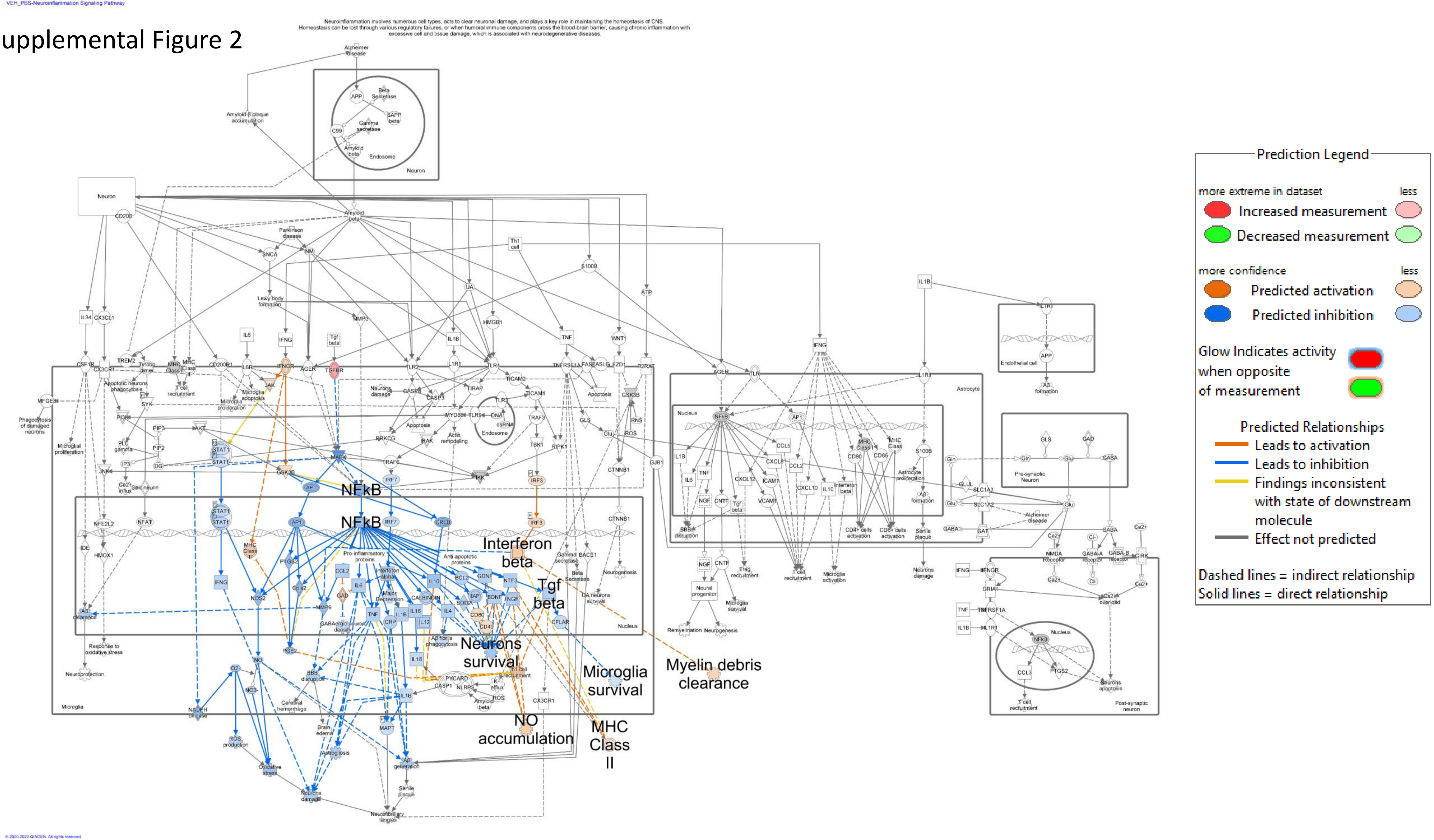

Neuroinflammation involves numerous cell types, acts to clear neuronal damage, and plays a key role in maintaining the homeostasis of CNS. Homeostasis can be lost through various regulatory failures, or when humoral immune components cross the blood-brain barrier, causing chronic inflammation with excessive cell and tissue damage, which is associated with neurodegenerative diseases.

### Supplemental Figure 3

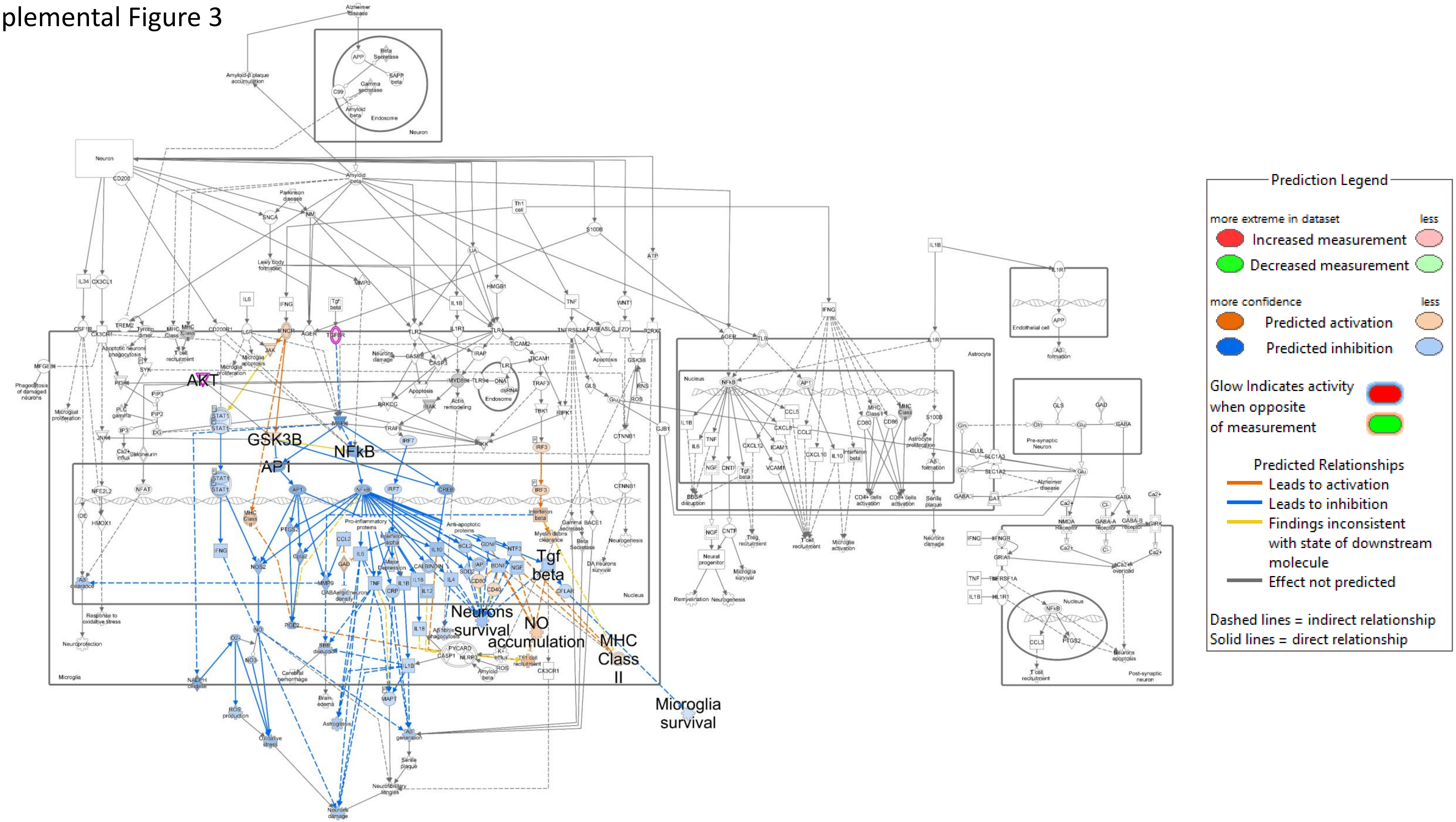


© 2000-2023 QIAGEN. All rights reserved.

### Supplemental Figure 4

Neuroinflammation involves numerous cell types, acts to clear neuronal damage, and plays a key role in maintaining the homeostasis of CNS. Homeostasis can be lost through various regulatory failures, or when humoral immune components cross the blood-brain barrier, causing chronic inflammation with excessive cell and tissue damage, which is associated with neurodegenerative diseases.

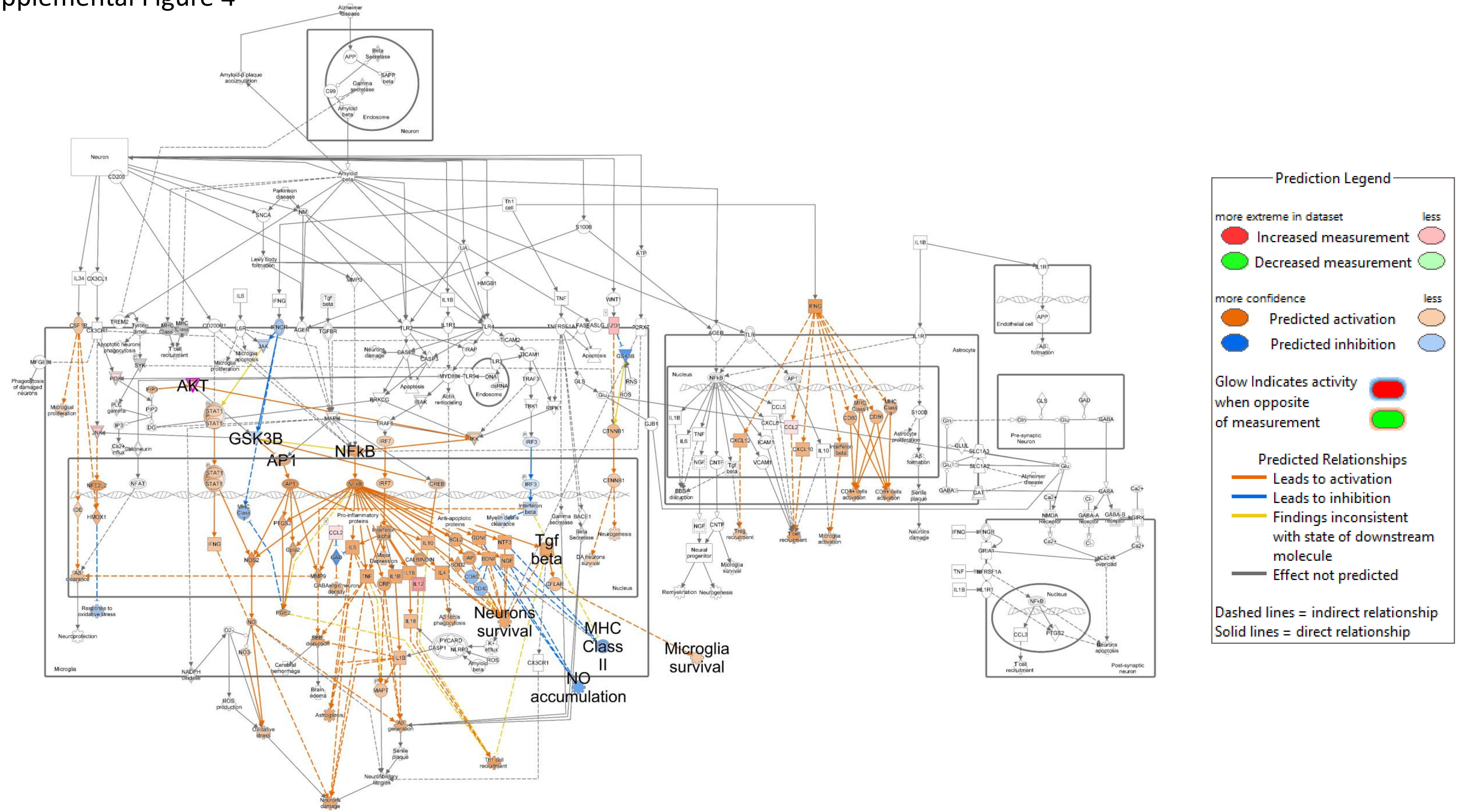


© 2000-2023 QIAGEN. All rights reserved.
