## Supplemental material - methods for "Extracellular condensates (ECs) are endogenous modulators of HIV transcription and latency reactivation"

**Running title: Extracellular condensates regulate HIV persistence**

**Methods:**

**Animal characteristics:** Eleven age- and weight-matched adult Indian RMs were randomly divided into 3 groups. Group-1 (n=4), [seven received twice daily injections of vehicle (VEH)] and were infected intravenously with 100TCID$_{50}$ of SIVmac251. Group-2 (n=4) received twice daily injections of THC similar to group 1 for four weeks prior to SIV infection until 6 months post SIV infection. Group 3 (n=3) served as controls and remained uninfected.

**Chronic administration of THC or VEH:** Administration of THC and Vehicle (VEH) was initiated by the intramuscular route four weeks before SIV infection at 0.18 mg/kg as used in previous studies [1, 2]. This dose of THC was found to eliminate responding in a complex operant behavioral task in almost all animals [3]. The dose was subsequently increased for each subject to 0.32 mg/kg, over a period of approximately two weeks when responding was no longer affected by 0.18 mg/kg daily (i.e., tolerance developed), and maintained for the duration of the study. The optimization of the THC administration in RMs accounts for the development of tolerance during the initial period of administration. Because in our previously published studies [1, 2] this dose of THC showed protection, the same dose was used in this study. The 0.32 mg/kg dose was also shown to be effective in SIV-infected RMs of Chinese origin [4]. SIV levels in plasma and BG were quantified using the TaqMan One-Step Real-time RT-qPCR assay that targeted the LTR gene [2, 5]. At necropsy, entire BG tissue was collected in RNAlater (Thermo Fisher Scientific) and Z-fix for total RNA extraction and embedding in paraffin blocks, respectively.

**Purification and characterization of basal ganglia derived EC:** The ECs used in this study were purified and characterized by Kaddour et al. 2023 [6].

**Cells lines:** The following reagents were obtained through the NIH HIV Reagent Program, NIAID, NIH: TZM-GFP Human Cell Line (JC.53 Derived), HRP-20041, contributed by David G. Russell and David W. Gludish [7]; HIV-1 Infected U937 Cells (U1), ARP-165 contributed by Folks et al., 1989 [8]; J-Lat GFP (ARP-9856) and J-Lat Tat-GFP (ARP-9851) cells were contributed by Dr. Eric Verdin [9, 10]. Hμglia (HC69) and CEMx174 cells were kind gifts from Dr. Jonathan Karn and Dr. Mahesh Mohan respectively. Hμglia (HC69) cells were maintained in Dulbecco's modified Eagle's medium (DMEM) (Gibco-BRL/Life Technologies) with low glucose, 1% fetal bovine serum (FBS), 1X glutamine, 50 ug/ml normocin and 1 μM of Dexamethasone. TZM-GFP cells were also maintained in DMEM but with 10% FBS. U1, J-Lat and CEMx174 cells were grown in RPMI (Gibco-BRL/Life Technologies) containing 5% exosome-depleted FBS (Gibco), 100 U/ml penicillin, 100 μg/ml streptomycin, sodium pyruvate, and 0.3 mg/ml l-glutamine (Invitrogen, Molecular Probes) as previously described by [11].

**EC internalization/Uptake:** PBS (control) or ECs were stained with SYTO™ RNASelect™ Green Fluorescent or DiR (DiIC18(7); 1,1′-dioctadecyl-3,3,3′,3′-tetramethylindotricarbocyanine iodide) red fluorescent using a final concentration of 5 μM of dye. PBS (control) or ECs were incubated for 20 minutes at 37°C and purified using

Exosome Spin Columns. Labelled ECs were added to cells for imaging using Lionheart FX automated scope. Images were acquired at 10X magnification at 0, 6 and 24 hours. Images were processed using Gen5 and raw data were plotted using GraphPad Prism 10.1 (La Jolla, CA, USA, www.graphpad.com).

**Real-time quantitative PCR (RT-qPCR):** 100,000 cells cultured overnight were treated with PBS or with 100 µg/mL of ECs for 04 days. Five µg total RNA was used for cDNA synthesis using the High-Capacity cDNA Reverse Transcription Kit (Applied Biosystems, Thermofisher). The cDNA was stored at −20 °C until use. The thermal cycler program and expression calculation was setup in a 7500 FAST real-time PCR system (Applied Biosystems, Thermofisher), and the fold change in gene expression was calculated using the standard ΔΔCT method as previously described [12-17].

**Western Blot Analysis of Viral Gene Expression:** A total of 50 µg of protein extracts from U1 cells treated with PBS, VEH, VEH|SIV, THC|SIV ECs for 4 days were subjected to 4–20% SDS-PAGE. The proteins were then transferred to a PVDF membrane, the membrane blocked with 5% BSA in 1× TBST buffer and incubated with relevant primary antibodies at 4 °C overnight. The blot was rinsed with 1× TBST 3–5 times for 5 min and incubated with relevant secondary antibodies for 1 h at room temperature. The blot was rinsed with 1× TBST 3–5 times for 5 min and images were captured using the Odyssey infrared imaging system (LI-COR). The integrated density of the band was measured using ImageJ 1.52a software.

**Viability Assay:** Cells (10,000 cells/well) were seeded in a 96-well plate. Cells were treated with VEH, VEH|SIV, THC|SIV ECs (50 µg/mL) or an equivalent volume of 0.1× PBS for 04 days at 37 °C. On day 04 of treatment, cells were counted, and the number of live cells determined using the Luna-II automated cell counter. In parallel, cell viability was determined using the trypan blue exclusion assay.

**RNA-Seq:** Isolated RNA (100-300 µg) sample quality was assessed by RNA Tapestation (Agilent Technologies Inc., California, USA) and quantified by Qubit 2.0 RNA HS assay (ThermoFisher, Massachusetts, USA). Paramagnetic beads coupled with oligo d(T)25 were combined with total RNA to isolate poly(A)+ transcripts based on NEBNext® Poly(A) mRNA Magnetic Isolation Module manual (New England BioLabs Inc., Massachusetts, USA). Prior to first strand synthesis, samples were randomly primed (5´ d(N6) 3´ [N=A,C,G,T]) and fragmented based on manufacturer's recommendations. The first strand is synthesized with the Protoscript II Reverse Transcriptase with a longer extension period, approximately 40 minutes at 42ºC. All remaining steps for library construction were used according to the NEBNext® Ultra™ II Directional RNA Library Prep Kit for Illumina® (New England BioLabs Inc., Massachusetts, USA). Final libraries quantity was assessed by Qubit 2.0 (ThermoFisher, Massachusetts, USA) and quality was assessed by TapeStation D1000 ScreenTape (Agilent Technologies Inc., California, USA). The final library size was about 430bp with an insert size of about 300bp. Illumina® 8-nt dual-indices were used. Equimolar pooling of libraries was performed based on quality control (QC) values and sequenced on an Illumina® Novaseq platform (Illumina, California, USA) with a read length configuration of 150 PE for 40M PE reads per sample (20M in each direction).

**Bioinformatics:** FastQC (version v0.12.1) was applied to check the quality of raw reads. Trimmomatic (version v0.39) was applied to cut adaptors and trim low-quality bases with default setting. STAR Aligner (version 2.7.10b) was used to align the reads. The package of Picard tools (version 3.0.0) was applied to mark duplicates of mapping. StringTie (version 2.2.1) was used to assemble the RNA-Seq alignments into potential transcripts. FeatureCounts (version 2.0.6) or HTSeq (version 2.0.3) was used to count mapped reads for genomic features such as genes, exons, promoter, gene bodies, genomic bins, and chromosomal locations. The TPM normalization method was used to report results because TPM results are sample independent, respects the average invariance, eliminates statistical biases inherent in other measure [18], and the TPMs are guaranteed to be the same across samples [19, 20]. Raw TPMs are provided in **Supplemental Table 1**.

**Secretome analysis using Cytokine Array/Immunoassay:** To determine the effect of ECs on the secretome of HIV latently infected cells, 100,000 U1 cells were counted and seeded in a 12 well plate with 500 ul of RPMI media. Cells were treated with 50 ug of VEH, VEH|SIV, THC|SIV ECs or 0.1X PBS for 04 days. Following the fourth day, cells were collected, triplicates were pooled and centrifuged at 500g for 5 minutes. Pellets were saved for other experiments (Transcriptome analysis) and supernatants were transferred to a fresh tube. Supernatants were clarified by centrifuging at 2000g for 10 minutes. Clarified supernatants were stored at -80 $^{o}$C for further analysis. 1 mL of supernatants from each group was used to perform cytokine array analysis following the manufacturer's protocol (Proteome Profiler Human XL Cytokine Array Kit Catalog Number ARY022B). https://resources.rndsystems.com/pdfs/datasheets/ary022b.pdf?v=20231215. Images of the membranes were acquired using Odyssey infrared imaging system Li-Cor CLX. Target protein expression as Dot blot were quantified using Empiria Studio ® Version 3.0 and protein expression presented as pixel intensity.

**Syncytia Formation Assay:** Syncytia assay was conducted using CEMx174 cells. Briefly, equivalent numbers (5000 cells/well) of CEMx174 cells were seeded in 48 well plate with 250 µl of RPMI media for 2 hours. After 2 hours, 250 µl of of clarified of supernatants from U1 cells treated with PBS, or 50 ug of VEH, VEH|SIV, THC|SIV ECs for 04 days were added to the cells. The cells were cultured at 37 °C. After 1, 2, and 4 days of culture, syncytia were visualized and imaged (10X) using Lionheart FX microscope.

**Cell to cell HIV transfer assay:** Cell-to-cell infection of TZM-GFP cells was performed with CEMx174 cells. Briefly, CEMx174 cells were labeled with PKHRed. The cells were with 1X DPBS to remove excess PKHRed label. Washed CEMx174 cells were cultured in the presence of supernatants from U1 cells treated with PBS or 50 ug of VEH, VEH|SIV, THC|SIV ECs for 04 days. Then, 1000 of the PKHRed-labeled CEMx174 cells were overlayed atop 1000 TZM-GFP cells. The co-culture was incubated for 02 days. Images of the cells were taken with LionHeart FX microscope at 10X magnification and analyzed with Gen5 3.10. The activation of GFP expression in TZM-GFP cells is a sign of virus transfer from CEMx174.

**Validation of transcriptome and secretome data:** Select genes from the transcriptome dataset were validated using total RNA isolated from independent experiments. The RNA samples were subjected to RT-qPCR and described above. On the other hand, validation of secretome data was conducted using ELISA [16]. Briefly, supernatants from HC69 Huglia cells were used to validate secretome cytokine array analysis. Briefly, 100,000 cells/well were plated overnight in a 12 well plate in 500 µl of DMEM-F12 supplemented with 1% FBS, and 1uM of Dexamethasone. The next day, cells were treated with VEH, VEH|SIV, THC|SIV ECs (50 µg/mL) or 0.1X PBS and incubated at 37°C for 04 days. Images were acquired each day to measure GFP intensity. At 04 day, supernatants were collected and clarified by centrifuging at 2000g for 10 minutes at 4°C and stored at -80 °C freezer for future use. To perform ELISA, high binding capacity 96 well plates (Costar 3166) were used. Multiple dilutions (1:1, 1:2, and 1:4) of HC69 supernatants (antigen) to coating buffer (100 mM of carbonate buffer) were made. Plates were coated with 100 µl/well antigen:coating buffer mix and incubated at 4°C overnight. For negative control, media:coating buffer (1:1) was used to coat the wells. Next day, plates were washed 3 times with ELISA wash buffer (Tris buffered saline with Tween 20, pH 8) and residual buffer removed by inverting the plate on paper towels. After washing, 100 µl of blocking buffer (5% BSA with NP-40) was added and plates were incubated at room temperature for 03 hours. After 03 hours, the plates were washed (as stated above), primary antibody (0.2 µg/mL) was added, and plates incubated at 4°C overnight. The next day, plates were washed and specie-specific HRP conjugated secondary antibody (1:10,000) was added, plates incubated at room temperature for 03 hours, washed, and excess media removed. 100 µl of TMB substrate (Abcam ab171523) was added to each well. After 15 minutes, stop solution (0.2M sulfuric acid) was added to each well. Absorbance was read at 450nm using Biotek synergy H1 reader.

**Validation of transcriptome and secretome data:** HC69 Huglia cells were used for data validation. Select genes from the transcriptome dataset were validated with RT-qPCR [21] using total RNA isolated from huglia cells, while secretome data were validated with supernatants for the huglia cells using ELISA [16].

**Flow Cytometry:** Equivalent numbers (100,000) of J-Lat -GFP or J J-Lat Tat-GFP cells were seeded in 48 well plate containing 500 µl of media. The cells were treated with 0.1X PBS or 50µg/mL VEH, VEH|SIV, THS|SIV ECs. Cells were cultured in the presence of PBS or ECs for 04 days. On day 04, cells were washed and resuspended in MACSQuant running buffer (Miltenyi Biotec) to perform flow cytometry using BD Flow cytometer. Analysis was performed using FlowJo™ v10.10.

**Statistical analysis:** Differential expression data were generated using Graphpad Prism. The significance cutoff was set to fold change (FC) >1.5 or <-1.5 and a p-value <0.05. Statistical differences were assessed by one-way ANOVA with Šídák's multiple comparisons test, two-way ANOVA with Tukey's multiple comparison test, or

Binary Student's t tests (Welch's correction) using GraphPad Prism 10. Details of specific statistics are presented in each figure legend where the p-values are listed, as well as in **Tables 2 to 4**.

**Abbreviations:**
BG – Basal ganglia
EC – Extracellular condensates
CNS – Central nervous system
THC – Delta-9-tetrahydrocannabinol
VEH – Vehicle
HIV – human immunodeficiency virus
SIV – simian immunodeficiency virus
